# Long-term forest-sector mitigation and radiative forcing under contrasting management, climate, and substitution pathways

**DOI:** 10.1101/2025.08.01.668130

**Authors:** I. Boukhris, F. Cherubini, A. Collalti, D. Dalmonech, C. Vonderach, G. Marano, F. Giannetti, S. Lahssini, M. Santini, R. Valentini

**Author notes:** francesco.cherubini.ntnu.no.

## Abstract

Forests are central to the EU’s climate neutrality strategy, currently offsetting ∼9% of total greenhouse gas emissions and offering further mitigation potential through harvested wood products and the substitution effect. However, the climate benefit of the forest sector is influenced by multiple interacting factors, including forest management, climate change, wood-use strategies, and assumptions about substitution benefits, as well as the timing and fate of carbon across the forest ecosystem and technosphere. To evaluate these drivers, we used a coupled forest growth and a wood products model to simulate five different silvicultural strategies under three climate change scenarios, four wood use schemes, and five displacement factor decay pathways over a 285-year period (2015–2300), applied to a *Pinus nigra* forest in Italy, questioning the impact of these factors on climate mitigation potential of the forest sector. We assessed forest sector balance (FSB, net carbon exchange between forest system and atmosphere), radiative forcing from biogenic CO_2_ (RF_bio_), and mitigation efficiency (ME) – the proportion of sequestered carbon contributing to net climate benefit. Results showed that FSB and RF_bio_ were “broadly” aligned, but ME varied with the magnitude and duration of biogenic emissions. The scenarios BIOE (bioenergy) and TM (modular cutting) achieved high FSB but showed lower ME due to concentrated or sustained emissions. WOOD (promotion of long-lived wood) and ADAPT (adaptation management) yielded higher ME under SSP1-2.6, while several strategies (WOOD, ADAPT, TRANS) became net sources under SSP5-8.5 after 2200. Substitution benefits declined under degressive assumptions, reducing mitigation by up to 53% especially for high-harvest scenarios. FSB was primarily shaped by climate and management, secondly by substitution, however; wood-use strategies had no significant long-term effect provided they did not impact resource availability. Together, these findings underscore that effective forest-sector mitigation requires not only maximizing cumulative carbon stocks, but also minimizing the magnitude, timing, and atmospheric residence time of emissions while carefully considering the role of substitution benefits.

**Highlights:**

- The long-term forest sector carbon balance is mainly governed by active management, climate conditions, and wood substitution pathways.
- Forest Sector Balance as a metric is “broadly” aligned with the radiative forcing from biogenic emissions
- The mitigation efficiency of forest sector options depends on emissions timing, duration, and amplitude
- Substitution factors also known as displacement factors need to be considered with greater caution

## 1. Introduction

Forests are key to the EU’s climate mitigation framework for achieving carbon neutrality by 2050. Currently they remove approximately 9% of total greenhouse gas (GHG) emissions from the atmosphere (EEA, 2022). Forest ecosystems serve as both carbon sinks, sequestering additional carbon, and reservoirs, storing it across several pools. When managed sustainably, forest carbon can be transferred laterally to harvested wood products (HWPs), establishing an external carbon pool and contributing to the substitution effect, as HWPs replace energy-intensive materials and fossil fuels, avoiding emissions (Rüter and Fulvio, 2025). In the EU, forests and HWPs together sequester approximately 400 Mt CO_2_-eq.yr^-1^, with HWPs contributing for about 10% to this total. Substitution provides a similar level of contribution with values ranging from 18 to 43 Mt CO_2_-eq.yr^-1^, further enhancing the forest sector’s overall mitigation potential (Jonsson et al., 2021; Nabuurs et al., 2017; Rüter S. et al., 2016).To assess forest sector mitigation, a system-wide perspective is needed to account for the distinct roles of carbon pools. This approach ensures that all relevant mitigation pathways, sequestration, storage, and substitution, are critically evaluated (Grassi et al., 2021; Lemprière et al., 2013). Forest-sector mitigation outcomes are shaped by multiple interacting factors. Understanding these drivers is essential to accurately assess the effectiveness and trade-offs among strategies. The forest carbon sink is primarily determined by both forest management and climate conditions (Ameray et al., 2021; Dye et al., 2024; Gregor et al., 2024; Liu et al., 2025). Similarly, the HWPs sink is affected by these factors but also depends on wood-use characteristics and the carbon footprint of the displaced material, which ultimately determines the substitution effect (Bozzolan et al., 2024; Brunet-Navarro et al., 2017; Budzinski et al., 2020). Beyond biophysical and technological factors, temporal aspects of carbon fluxes also play a critical role.

Among these factors, forest management has received particular attention (Valentini and Miglietta, 2015). Practices range from passive conservation to intensive silvicultural interventions (Barredo et al., 2024; Dalmonech et al., 2022; Pilli et al., 2022). An ongoing debate in the EU revolves on whether active or reduced management contributes more effectively to climate mitigation (Luyssaert et al., 2007; Noormets et al., 2015). In managed forests, early interventions such as tending and thinning are used to improve wood quality, promote tree growth, and reduce the likelihood of natural mortality. As a result, more wood can be harvested for use in HWPs instead of being lost to natural decay. At the same time, these operations enhance forest resilience to disturbances, which are becoming more frequent due to climate change (Barrette et al., 2023; Kilpeläinen and Peltola, 2022; Moreau et al., 2022). In the long term, while unmanaged forests tend to maintain higher carbon stocks, especially in the dead wood pool (Nagel et al., 2023), studies suggest that managed forests can achieve a superior overall carbon balance when accounting for the substitution effect (Krug et al., 2012; Schulze et al., 2020).

Differences in the mitigation potential of multiple active management strategies exist and are primarily driven by factors such as structure complexity, rotation length, thinning intensity, and thinning intervals. In parallel, at least two key factors influence the HWPs sink: the recycling rate and the lifespan of HWPs, both of which delay carbon emissions to the atmosphere through combustion or decomposition in disposal sites (Brunet-Navarro et al., 2017; Pingoud et al., 2003; Wei et al., 2023). Finally, one of the most critical assumptions in estimating the forest sector’s substitution effect is the assumption of stationary displacement factors (DF). Assuming fixed DF fails to account for potential improvements in energy efficiency and reductions in the carbon footprint of competing sectors (e.g., steel, concrete, and energy) that may lead to reductions in GHG emissions. Any advancements in these sectors would consequently diminish the substitution effect of wood (Brunet-Navarro et al., 2021; Harmon, 2019; Howard et al., 2021). Therefore, assessing alternative strategies with a fixed DF reflects only the maximum substitution potential, while the actual effect is likely lower and often overestimated in global reports.

A critical yet often overlooked aspect of climate mitigation assessments is the timing of carbon fluxes and how this aligns with the time horizon of mitigation projects. As Levasseur (2010) highlighted, “*carbon neutrality does not equate climate neutrality*”, as carbon persists in the atmosphere before being reabsorbed, contributing in the meantime to the climate forcing (Levasseur et al., 2010). Forest-Technosphere systems that release carbon rapidly but sequester it slowly can have a stronger short-term climate impact than those with faster sequestration rates and slower carbon release (Holtsmark, 2012; Laganière et al., 2017). These short-term emissions influence radiative forcing, potentially intensifying near-term warming and affecting long-term climate stability. Similarly, the time horizon, the timeframe over which forest systems are evaluated, significantly shapes their assessed climate impact. Over short time frames, immediate emissions from harvesting may outweigh sequestration benefits, resulting in a higher net climate impact. In contrast, longer horizons allow forest regrowth along with the other climate system components to offset initial emissions, improving mitigation outcomes (Cherubini et al., 2011; Holtsmark, 2013; Sierra et al., 2021). Given this, assessing the mitigation potential of forests based on carbon balances alone is insufficient; it must also account for their contribution to radiative forcing over time. To capture these temporal dynamics, we first quantified climate impact using a timing-sensitive indicator, RF_bio_, which integrates the net carbon fluxes over time through a global impulse response function. However, while RF_bio_ describes the magnitude of climate impact, it does not indicate how efficiently carbon is processed to achieve that outcome. To address this limitation, we introduce the concept of mitigation efficiency (ME), a metric reflecting the climate benefit delivered per unit of carbon cycled through the forest sector. Together, RF_bio_ and ME offer a more nuanced basis for evaluating forest-sector strategies, emphasizing not just how much carbon is removed, but how effectively and durably that removal contributes to climate mitigation.

Despite the well-established role of forests and HWPs in climate mitigation, significant uncertainties remain regarding the long-term effectiveness of different forest management strategies. These uncertainties arise from the complex interactions among forest growth, harvest intensity, product use, and substitution dynamics, as well as from simplifying assumptions, such as stationary DF. Moreover, most studies focus on short-term carbon balances, overlooking the long temporal cycles of forest systems and the fact that mitigation potential may emerge at different timescales depending on the applied management (Metzler et al., 2024; Skytt et al., 2021).

The main aim of this study is to quantify the long-term climate-mitigation performance of the forest sector by jointly accounting for forest carbon dynamics, HWPs, and time-dependent substitution effects under climate change. To achieve this, we compare five forest management strategies under three climate change scenarios over a 285-year period, evaluate how alternative displacement pathways and the decay of displacement factors influence mitigation outcomes, assess whether high carbon balances consistently correspond to stronger climate benefits across different time horizons, and determine how the mitigation efficiency of the forest-sector options emerges and evolves over time.

We addressed these questions using an integrated modeling framework coupling a process-based forest model with a wood product model (WPM) in a factorial experimental design. The framework was applied to a *Pinus nigra* J.F. Arnold stand in the Vallombrosa Forest (Tuscany, Italy), representative of temperate European forests. *P. nigra* was selected due to its wide ecological adaptability, increasing use in European reforestation programs, and relevance for climate-resilient forestry (EU council, 2025; Vacek et al., 2023).

## 2. Materials and Methods

### 2.1. Study area

The study was conducted in the state forest of Vallombrosa (43° 44’ N, 11° 34’), a biogenetic reserve of approximately 1000 ha in Tuscany, Italy. The forest experiences a mean annual precipitation of about 1195 mm and a mean annual temperature slightly higher than 10 °C. January is the coldest month (1.9 °C), July the warmest (19.6 °C). All the conifer plantations were traditionally managed as even-aged. *Pinus nigra* was introduced in the forest (Bottalico et al., 2012) as part of the main reforestation program that occurred after the second World War. These stands cover 11% of the total forest area, with ages ranging from 60 to 95 years. From 2005, as reported in the Vallombrosa forest management plan (*Piano di gestione e silvomuseo*) (Travaglini, 2009). *Pinus nigra* stands are managed under close-to-nature silviculture principles following the systemic silviculture approach (Ciancio and Nocentini, 2011), with the principles of modular/selective continuous cutting. This approach rooted in Continuous Cover Forestry (CCF) and prescribed by Ciancio (2004) promotes resource sustainability and natural regeneration by dynamically reassessing the harvestable volume (Ciancio and Nocentini, 2004). The silvicultural systemic method for growing stock harvesting calculates the real wood provision over a 5-year period. Thinning is allowed only if the remaining standing volume is kept above the minimum threshold (typically 100-150 m^3^ ha^-1^ for *Pinus nigra*). If the condition is not met, only the portion of increment that preserves the minimum stock is thinned. Thinning is also carried out to remove trees at high risk of falling. When harvesting is conducted, the resulting residues are left on site. To monitor forest yield and health, and in the context of the forest management plan, local permanent forest inventory plots were established. The permanent circular plots (530 m²) are identical to those of the Italian National Forest Inventory (NFI; https://www.inventarioforestale.org) and were surveyed in 2005. Several of these plots are located within *Pinus nigra* stands and were resurveyed again in 2017, 2018, and 2023.

### 2.2. Study key silvicultural and modeling assumptions

The following assumptions are considered in this study: the selected forest stand is assumed to be representative of the average conditions across the broader *Pinus nigra* landscape. The same forest management practices and wood use behavior are consistently applied/maintained throughout the study period. Additionally, species composition is assumed to remain unchanged, meaning no species conversion occurs over the time horizon. Finally, each management scenario assumes a starting point of a recent plantation established in 2015 following a clear-cut, with a planting density of 2500 seedlings per hectare. Tree dimensions at establishment followed species-specific default parameters of the growth model (initial height = 1.3 m). Biomass and soil carbon pools were initialized using a 2500-year spin-up to equilibrium under local climate conditions prior to the start of simulations. All management scenarios began from identical stand conditions, ensuring that the differences among scenarios arise solely from management, climate forcing, and wood-use assumptions.

In addition, landfilling activities were assumed to gradually phase out by the year 2060, following an exponential decay model calibrated on national data. This estimate is based on the observed reduction in the share of Italian municipalities utilizing landfill, which declined from 46% in 2010 to 18% in 2022 (EEA, 2024). By assuming a constant relative rate of decrease, landfilling is projected to approach zero use by 2060, and this cutoff year is adopted as the upper limit for landfill emissions modeling.

### 2.3. Definition of scenarios

#### 2.3.1. Climate change pathways

We investigated two coupled representative concentration pathways (RCPs) and shared socioeconomic pathways (SSPs) based on distinct emission scenarios to assess potential future climate trajectories and their socioeconomic implications. The SSP1-RCP2.6 scenario (SSP1-2.6 hereafter) represents a world of sustainability-focused growth and equality, and it assumes that strong climate protection measures are implemented, aiming to limit global warming to well below 2°C above pre-industrial levels (O’Neill et al., 2014; Riahi et al., 2017; van Vuuren et al., 2011). In contrast, SSP5-RCP8.5 (SSP5-8.5 hereafter) reflects a world of rapid and unconstrained growth in economic output, energy demand and use, as well as limited to no mitigation actions and a persistent reliance on fossil fuels (Calvin et al., 2017; Meinshausen et al., 2011; Rogelj et al., 2018). Climate projections were obtained from the Coupled Model Intercomparison (CMIP 6) project (Eyring et al., 2016), covering the period from 2015 to 2300. Data were sourced from the MRI-ESM2-0 global climate model, as it was the only model available that simulated the variables of interest over the long term, extending to the year 2300. The outputs were downscaled to a 0.25° resolution using the perfect prognosis downscaling method. A set of statistical models and algorithms was initially calibrated using historical (observed) data from both coarse predictors (ERA5 reanalysis; (Hersbach et al., 2020) and local predictand data (E-OBS reanalysis, including precipitation, maximum, and minimum temperature; Cornes et al., 2018) over a representative 30-year climate period (1991–2020). The methods were then validated and evaluated, with the best-performing models selected to downscale daily precipitation, maximum, and minimum daily temperatures. Moreover, atmospheric CO_2_ concentrations for different climate trajectories were obtained from (Meinshausen et al., 2020). Additionally, we simulated the current climate (CUR) for the period 2020–2300 by randomly sampling each day of the year from the local station records (Vallombrosa, locality of Reggello) over the period 2005–2023 (see **Figure 1**).

**Figure 1:**
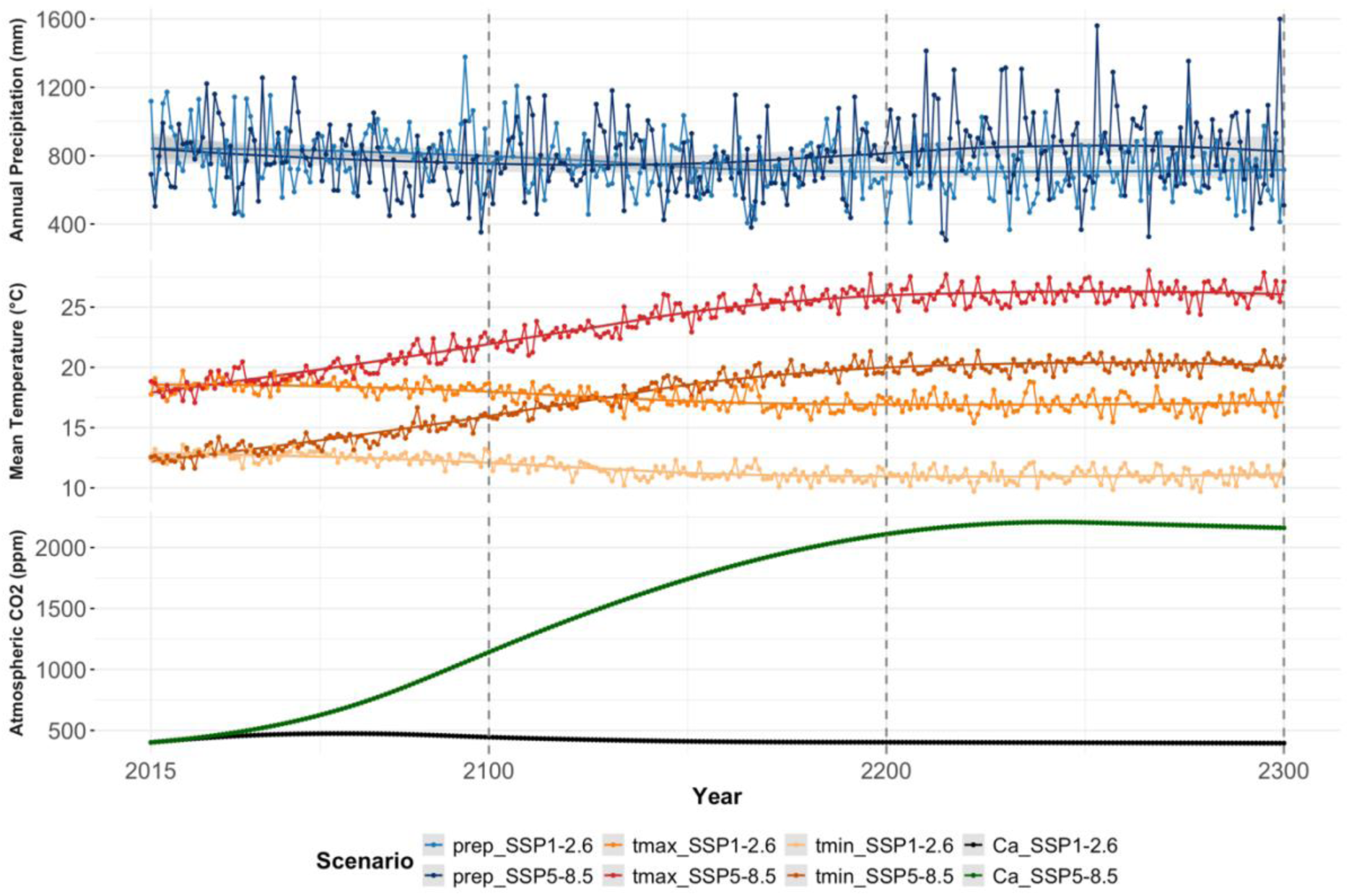
Climate projections from the downscaled MRI-ESM2-0 model under the SSP1-2.6 and SSP5-8.5 scenarios, driving the simulation for the period 2015–2300. The top panel shows annual precipitation (mm), the middle panel displays the annual mean of daily maximum and minimum temperatures (°C), and the bottom panel shows atmospheric CO₂ concentration (ppm), labeled as Ca.

#### 2.3.2. Forest management scenarios

Five management scenarios were developed, grounded in five policies outlined in the EU Forest Strategy (European Commision, 2021), the EU Bioeconomy Strategy (European Commision, 2018), and the EU Forest Biodiversity Strategy (European Commission, 2021). These scenarios were designed to reflect the specific characteristics of the target species and the study area. In this process, the regulations for *P. nigra* management in the Tuscany region were also considered (refer to (Cantiani P et al., 2018) for further details). Each policy was subsequently translated into stand-based forest management plans with the expert input of local stakeholders, including forest managers and academic scholars.

The silvicultural interventions applied across the scenarios include pre-thinning and thinning operations, implemented with varying intensities and intervals. In shelterwood-based scenarios, additional interventions such as preparatory cuts and seedling cuts are employed to facilitate natural regeneration under partial canopy cover. Final cuts are applied in both shelterwood and clearcut systems, serving to remove the remaining overstory once regeneration is established in shelterwood, or to harvest the entire stand at once in clearcutting scenarios. In the latter case, artificial regeneration follows harvest operations, with planting carried out at a density of 2500 seedlings per hectare (See **Figure 2** for detailed silvicultural intervention schedules).

**Figure 2:**
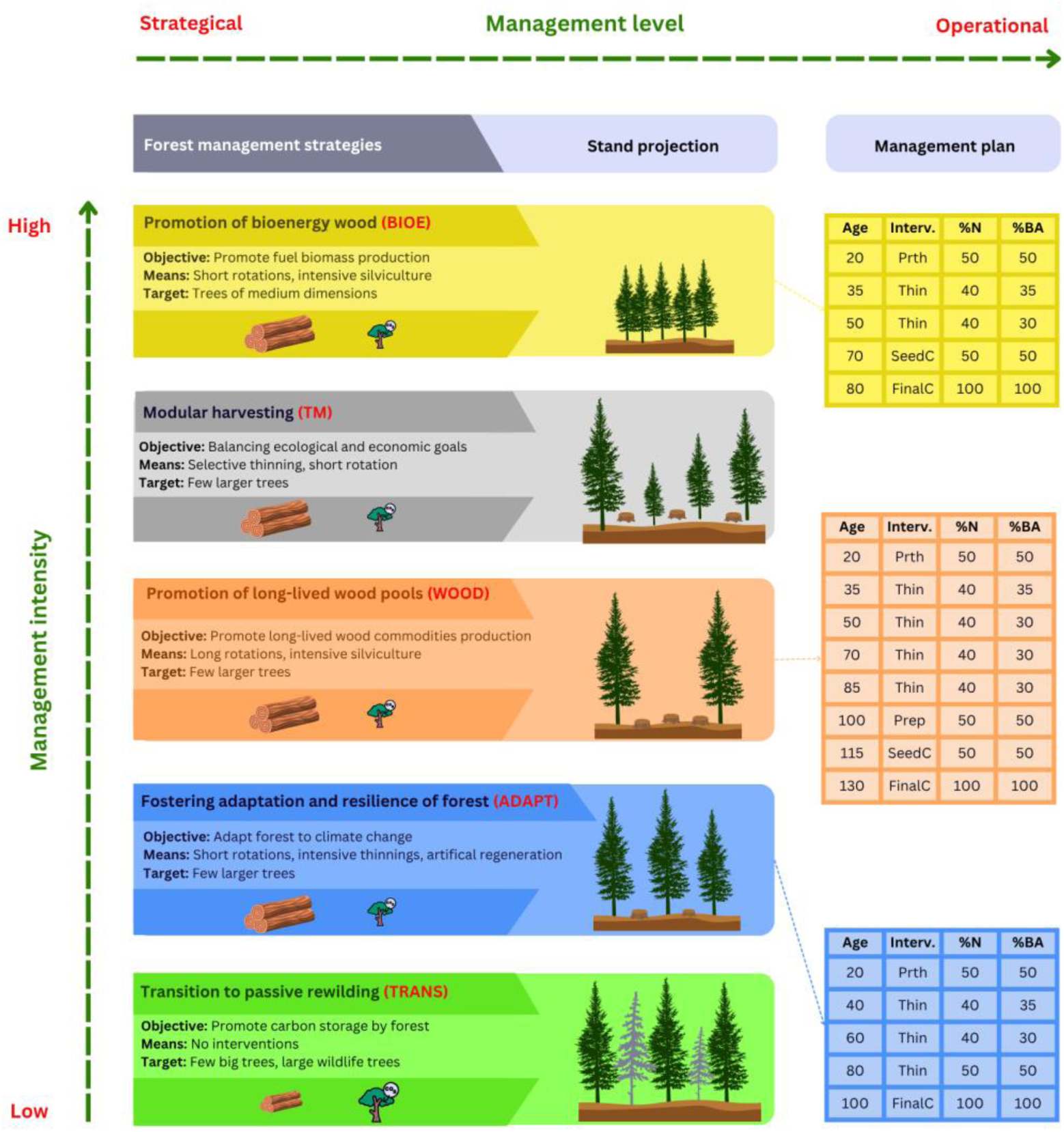
Overview of the five forest management strategies evaluated in this study, arranged along a gradient of management intensity from high (top) to low (bottom). Strategies include: BIOE, TM, WOOD, ADAPT, and TRANS. Each strategy is defined by its objective, silvicultural approach, and target stand characteristics, progressing from strategic objectives to operational implementation. The right-hand panels show the corresponding management plans for each strategy, including timing and type of intervention (i.e., Prth: pre-thinning, Thin: thinning, Prep: preparatory cut, SeedC: Seeding cut, and FinalC: final cut), percentage of trees removed (%N), percentage of basal area removed (%BA) scheduled at specific ages.

The first scenario is known as ‘*Tagli Modulari’* which represents modular cutting (hereafter TM), and reflecting the typical practices applied in the Vallombrosa forest and described in detail in Section 1. The “Wood-Based Bioenergy Production” (BIOE) assumes that the EU is transitioning to a more sustainable energy future by gradually promoting the use of wood bioenergy as a replacement for fossil fuel energy. While direct use of wood logs for energy is discouraged under the EU’s Renewable Energy Directive (RED II and RED III), only the portion of trees unsuitable for higher-value uses are directly allocated to bioenergy. Practically, this management plan consists of a short rotation (70-80 yrs) and the application of 1-3 thinning treatments. The “Promotion of long-lived HWPs” (WOOD) explores how forest management can support climate mitigation by increasing the supply of high-quality timber suitable for durable HWPs. The scenario assumes that domestic wood use in the EU shifts increasingly toward long-lived applications such as construction materials, which can store carbon for extended periods and substitute for more carbon-intensive materials. While the lifespan of HWPs also depends on post-harvest factors, such as recycling systems and end-use, this scenario emphasizes the forest management component that facilitates the availability of timber suitable for such uses. On the forest side, this objective is supported by extending rotation lengths (e.g., 120-150 yrs) aiming to harvest trees with larger diameters and higher wood quality through stronger thinning. The “Enhancing Forest Adaptation and Resilience” (ADAPT) aims to strengthen the resilience of forests, particularly Mediterranean forests, which are already experiencing severe drought conditions. Practically, this management plan will consist of applying intensive thinning to alleviate competition on site resources and adopting short rotation cycles to alleviate disturbance risks, in addition to promoting artificial regeneration. The last scenario, “Transition to passive rewilding” (TRANS) is designed to facilitate the shift from structured stand management toward a self-regulating forest system with minimal long-term disturbance. Initial interventions are intended to accelerate natural processes such as gap creation, early mortality, and canopy opening, which commonly occur in the late stages of forest succession and structural diversification (Bauhus et al., 2009). These actions aim to promote structural diversification and regeneration. Thinning is applied at regular intervals between 2030 and 2150, with intensities ranging from 30% to 50% of the stand basal area. After 2150, no further interventions were applied, allowing the forest to develop autonomously under passive conditions.

#### 2.3.3. Wood use scenarios

To explore the potential implications of improved cascading through an increase in product lifespan and recycling rates, four different wood use scenarios were developed. The first, the BAU as “Baseline Scenario”, maintained constant recycling rate and lifespan values. In the LS20 as “Longevity Scenario”, the lifespan of products was increased by 20%. The RR20 as “Reuse Scenario” focused on increasing the recycling rate of products by 20%. Finally, the RRLS20 “Sustainability Scenario” increased both the lifespan of products and the recycling rate by 20%. In BAU, the recycling rate of wood products was fixed at 10% through the simulation period while product lifespans were parametrized from published studies (please refer to **Table S1.5**).

#### 2.3.4. Substitution scenarios

In addition to a baseline scenario (SUB0), which assumes that the DF remains constant over time, we considered four decarbonization pathways with varying linear rates of DF reduction. We adopt either a gradual (25% DF reduction by 2050, called SUB25), moderate (50%, SUB50), significant (75%, SUB75) or a rapid (100%, SUB100), the latter being in line with the EU’s climate neutrality targets. DF values were parametrized based on literature sources (Sathre and O’Connor, 2010; Suter et al., 2017) (please refer to Table S1.6).

### 2.4. Modeling framework

#### 2.4.1. Modelling system and key concepts

The system considered in this study includes both forest and HWPs pools, collectively representing the forest sector. It accounts for incoming and outgoing carbon fluxes while disregarding internal exchanges, as these remain within the system and do not affect its net balance. The forest sector system consists of five key components: (1) Aboveground biomass, comprising stem wood, branches, stumps, and foliage of live trees, where the focus is on CO_2_ fixation through photosynthesis and biomass accumulation; (2) Belowground biomass, including fine and coarse roots of live trees, also assessed for CO_2_ fixation; (3) Aboveground Dead Organic Matter (ADOM), consisting of standing dead trees, downed woody debris, litter, and humus, where carbon is gradually released through decomposition; (4) Belowground Dead Organic Matter (BDOM), encompassing soil organic matter and dead roots, with carbon emissions primarily occurring through microbial respiration; and (5) HWPs, representing carbon stored in HWPs during their use phase, with CO_2_ emissions occurring over time through disposal in landfills or combustion for bioenergy (see **Figure 3**). Beyond these carbon pools, the substitution effect plays a critical role in the forest sector’s overall climate mitigation. The annual forest ecosystem carbon balance (FEB; tC ha^-1^ yr^-1^) is defined as:

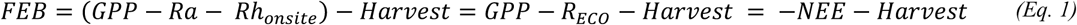

where GPP represents the gross primary production; *Ra*, *Rh*_*onsite*_, and *R*_*ECO*_ correspond to autotrophic, heterotrophic, and ecosystem respirations occurring within the forest, respectively, Harvest accounts for biomass removal through forestry operations, and NEE represents the net ecosystem exchange within the forest. All terms are expressed in units of tC ha^-1^ yr^-1^.

**Figure 3:**
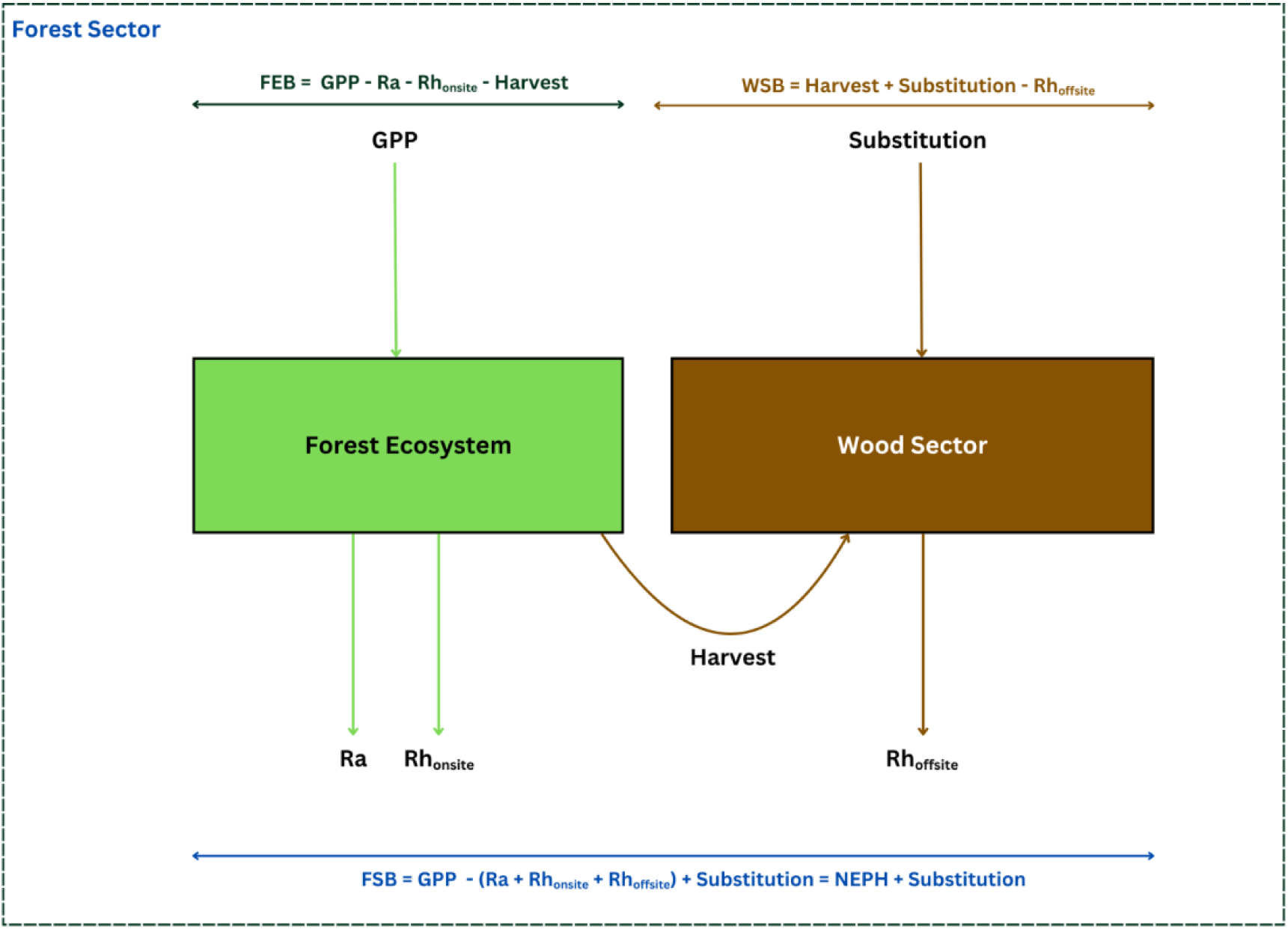
Conceptual representation of carbon flows and balances with the forest sector, divided into the forest ecosystem (green box) and wood sector (brown box). Gross Primary Productivity (GPP) constitutes the carbon inflow to the forest ecosystem while both Ra and Rh_onsite_ constitute the carbon outflow from the forest ecosystem. Harvested biomass is transferred from the forest ecosystem to the wood sector where it contributes to substitution benefits and eventually to Rh_offsite_.

Harvested biomass enters the wood sector, where it is temporarily stored in HWPs before eventually being released as emissions through disposal, or combustion for bioenergy. Therefore, the wood sector carbon balance (WSB; tC ha^-1^ yr^-1^) is expressed as:

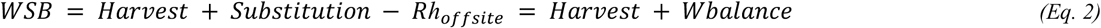

where *Substitution* represents avoided emissions resulting from the replacement of fossil fuels and energy-intensive materials with HWPs, *R*h**_*offsite*_ (tC ha^-1^ yr^-1^) refers to the heterotrophic respiration occurring outside the forest, mainly from HWPs decomposition and combustion, and *Wbalance* refers to the balance between *Substitution* and *R*h**_*offsite*_, both of which are expressed in units of tC ha^-1^ yr^-1^.

The annual overall forest sector balance (FSB; tC ha^-1^ yr^-1^) integrates both forest and wood sector dynamics, incorporating the mitigation benefits of substitution:

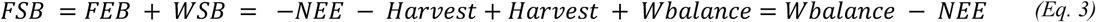

The terms *R*h**_*onsite*_, and *R*h**_*offsite*_ follow the terminology established by (Schulze et al., 2021).

#### 2.4.2. Modelling framework and simulation setup

A forest sector modeling framework was developed by coupling 3D-CMCC-FEM (v5.7 BGC) (Collalti et al., 2014; Dalmonech et al., 2024; Puchi et al., 2026; Saponaro et al., 2025; Vangi et al., 2026), a stand-level, climate-sensitive, process-based model, with TimberTracer (Boukhris et al., 2025) a WPM that tracks carbon stocks in HWPs, substitution effects, and emissions from disposal sites (see **Figure 4**). The choice of 3D-CMCC-FEM was motivated by its suitability for long-term climate-mitigation analyses. The model explicitly represents photosynthesis, respiration, and carbon allocation, allowing sensitivity to changing climate conditions and capturing climate-carbon feedback affecting growth and mortality. Its process-based stress-response formulation enables realistic simulation of drought induced decline, while the structural representation of stand dynamics provides a mechanistic description of mortality and competition at the species scale. At the same time, the model remains computationally efficient, which is essential for factorial multi-century simulations and sensitivity analyses. Its relatively transparent parametrization further facilitates uncertainty assessment and interpretation. Complementarily, TimberTracer was selected to represent the technosphere component of the forest sector. Based on a material-flow approach, the model simulates the temporal dynamics of carbon storage and emissions across all HWP pools and disposal pathways, while explicitly accounting for material and energy substitution effects. Its compatibility stand-level inputs allow seamless coupling with forest growth models such as 3D-CMCC-FEM, enabling consistent carbon accounting from forest ecosystem processes to post-harvest product use. The modular structure of the model further supports scenario analysis and sensitivity testing, making it suitable for evaluating long-term mitigation pathways.

**Figure 4:**
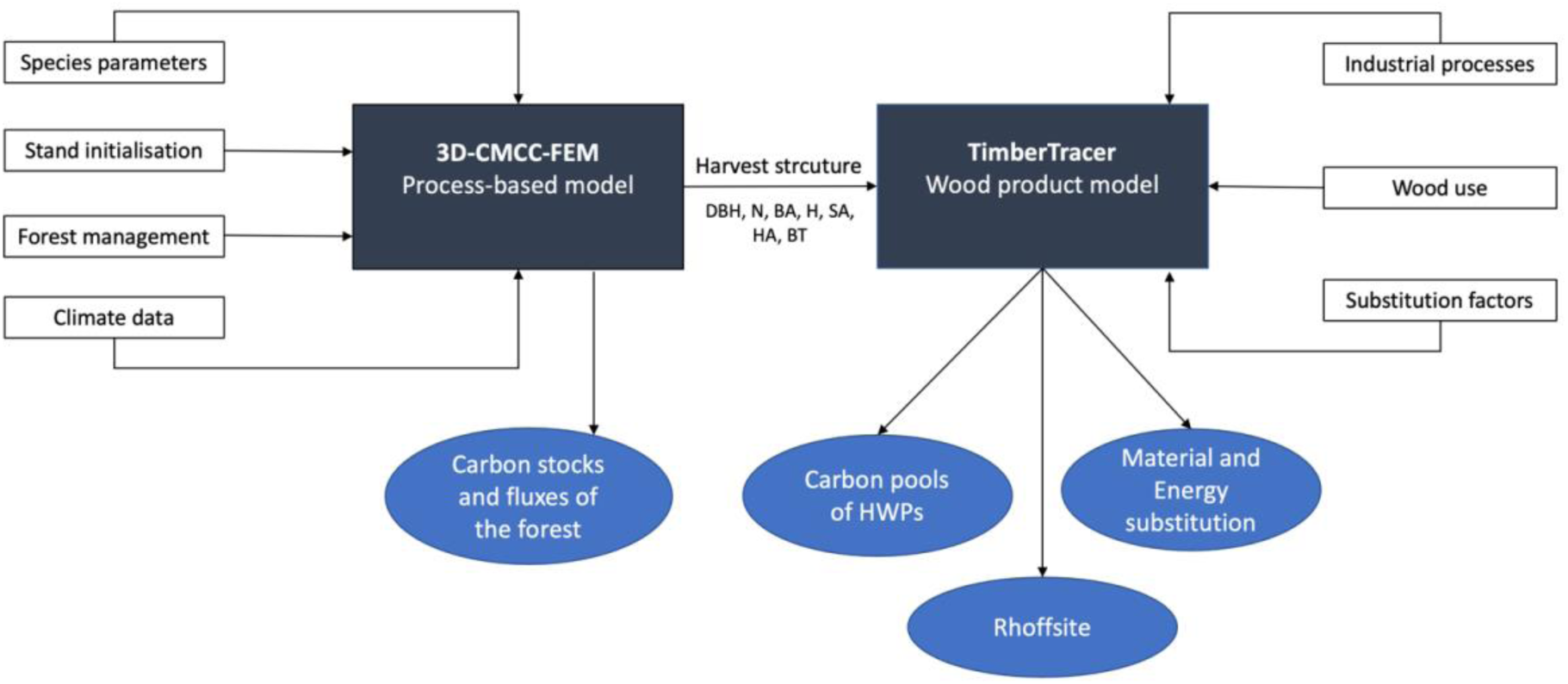
Carbon modeling framework coupling a forest process-based model (3D-CMCC-FEM) and TimberTracer, a WPM. DBH is the mean diameter at breast height, N is the stand density (ha^-1^), BA is the basal area (m^2^ ha^-1^), H is the mean tree height (m), SA is sapwood area of the mean tree (cm^2^), HA is the heartwood area of the mean tree (cm^2^), and BT is the bark thickness of the mean tree (cm).

Given the complexity and level of detail associated with each model, including their structure, parametrization, input data, and validation procedures, full technical descriptions have been moved to the Supplementary Materials (**Tables S1.1-S1.7**) to maintain clarity and focus on the main text.

#### 2.4.3. Modeling the climate impact of the forest sector

To assess the climate impact of the forest sector, we model atmospheric CO_2_ emissions decay using an Impulse Response Function (IRF), which quantifies the temporal reaction of a dynamic system in response to some external change (Joos et al., 2013). The multi-model mean from the carbon cycle model intercomparison in Joos et al. (2013) is used to simulate this decay process, incorporating a multi-exponential decay function to simulate the response of different carbon pools, accurately reflecting the transient and long-term behavior of carbon in the atmosphere, land, and ocean reservoirs (Cherubini et al., 2011; Joos et al., 2013; Strassmann and Joos, 2018). Its associated IRF is mathematically expressed as:

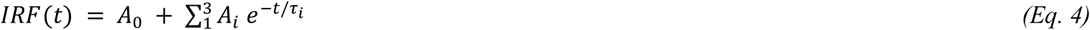

where *A_0_* = 0.2173, *A_1_*= 0.2240, *A_2_*= 0.2824, *A_3_*= 0.2763, *τ_1_* = 394.4, *τ_2_* = 36.54, *τ_3_* = 4.304 The amplitude *A_0_* represents the asymptotic airborne fraction of CO_2_, which remains in the atmosphere in the long term due to the equilibrium response of the ocean–atmosphere system. Each *A*_*i*_ is associated with a characteristic residence time *τ*_*i*_, which describes how long the corresponding fraction of carbon remains in the atmosphere before being taken up by the combined action of terrestrial and oceanic processes. Together, the coefficients (*A_0_* + *A*_1_ + *A*_2_ + *A*_3_) sum to 1, and thus each *A*_*i*_ can be interpreted as the fractional contribution of a given decay component to the overall atmospheric response. It is important to note that these components do not correspond to individual, physically distinct carbon sinks, but rather to a mathematical approximation of the combined response of the global carbon cycle.

The atmospheric concentration of CO_2_ over time can be described as the convolution of the IRF with the net carbon flux. It is expressed as the integral of past net emissions at time t’, weighted by the fraction of CO_2_ remaining in the atmosphere after a time interval t -t’, where t is the reference time point.

To compare the climate impact of biogenic CO_2_ emissions across different forest-sector scenarios, a consistent metric is required. In this paper, we developed an approach based on the integrated radiative forcing concept adapted for biological systems, referred to here as RF_bio_. While originally formulated for pulse emissions in Life Cycle Assessment contexts (e.g., GWP_bio_; see Cherubini et al., 2011), here we extend the concept to account for net carbon fluxes over time, making it applicable to dynamic forest-sector modeling. RF_bio_ is derived from the time-integrated radiative forcing caused by emissions, quantifying the perturbation of the Earth’s energy balance. It is calculated as follows:

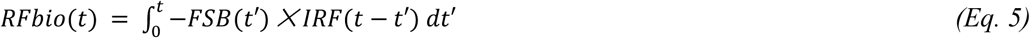

A positive RF_bio_ indicates a net warming effect. In contrast, an RF_bio_ equals to or less than zero implies that the system has no net contribution to atmospheric warming. This condition can therefore be interpreted as the forest-sector system being climate neutral (RF_bio_ ≈ *0*), adds to climate warming (RF_bio_ > 0), or climate cooling (RF_bio_ < 0) over the assessed time horizon. Figure 5 provides an illustration of these concepts.

**Figure 5:**
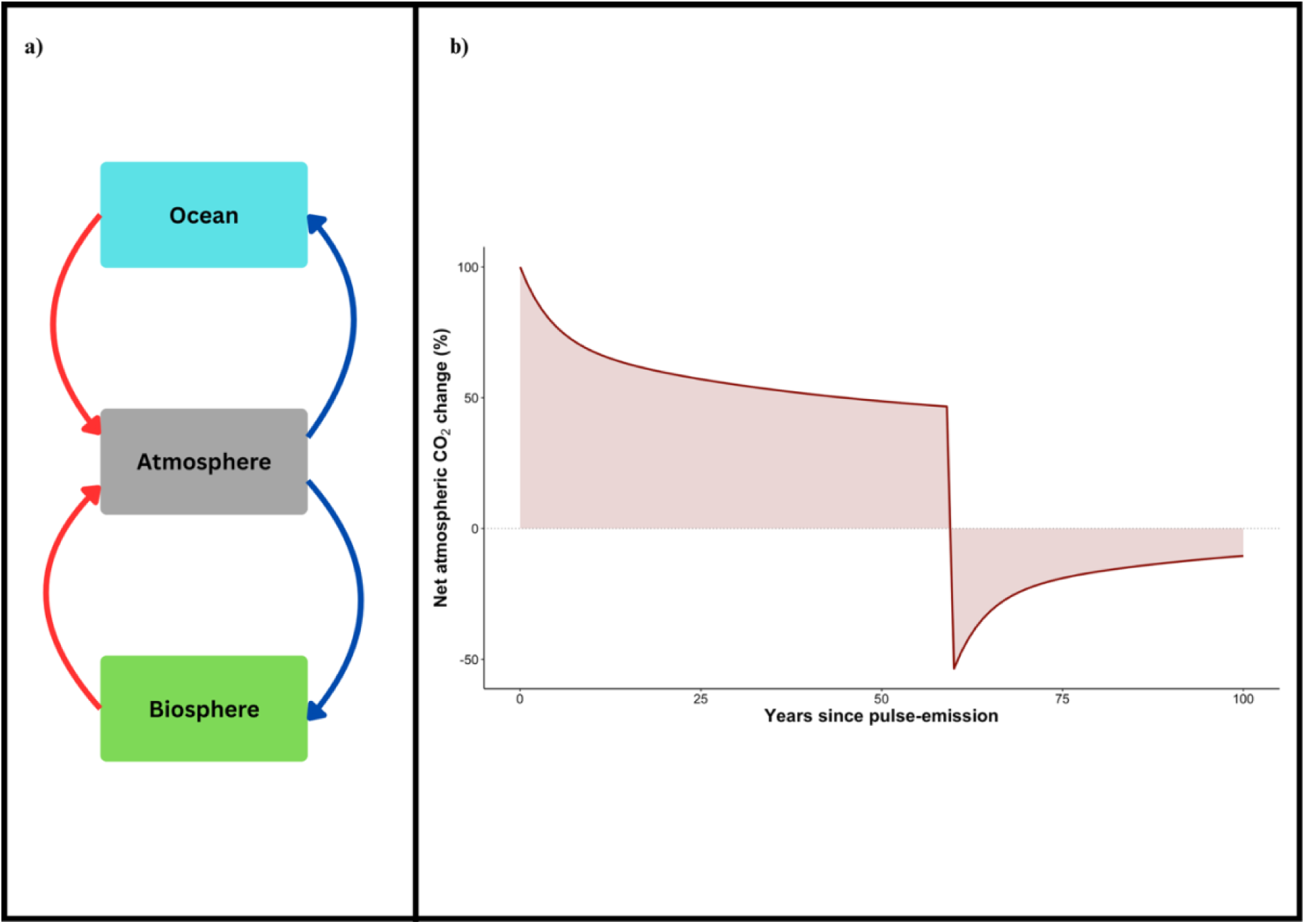
CO_2_ dynamics across Earth system compartments and the carbon removal dynamics. (a) Schematic representation of CO_2_ exchanges between the atmosphere, ocean and biosphere. Red arrows indicate CO_2_ emissions to the atmosphere; blue arrows indicate CO_2_ uptake from the atmosphere. (b) Net atmospheric CO_2_ change (%) over time following a pulse emission, with equivalent carbon removal occurring at year 60. For analogy, assume that by the starting year, harvested wood is burned for bioenergy and the resulting emissions are fully captured by year 60. Response is simulated using the IRF Bern 2.5 CC model. The shaded area represents the RF_bio_, highlighting temporary overshoot and long-term legacy effect. Positive values, denote a forcing effect while negative values denote a cooling effect.

The modeling approach described in Sections 4.2 and 4.3 is operationalized through an integrated methodological workflow, illustrated in Figure A.1.

### 2.5. Assessment of carbon and climate mitigation potentials

Carbon balance (CB) was estimated by tracking net changes in forest carbon pools and HWPs, while also accounting for the generated substitution effects. CB was determined by summing the FSB (see Eq. 3) over a time horizon T:

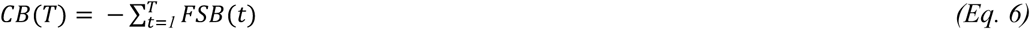

where *FSB*(*t*) represents the annual forest sector balance at time t. The total time horizon T was considered over 85, 185, or 285 years. The minus sign in Equation 6 reflects the convention that a negative CB indicates a net carbon sink (i.e., net removal of CO_2_ from the atmosphere), while a positive value indicates a net source (i.e., net emissions).

Climate mitigation potential, expressed as radiative forcing due to forest management, was assessed using the RF_bio_ metric (see Eq. 5), which accounts for the temporary nature of biogenic CO_2_ emissions and their atmospheric decay over time.

Furthermore, the concept of ME is introduced to measure the effectiveness of mitigation efforts, is defined as the ratio between the negative RF_bio_ and the corresponding carbon balance. The negative sign ensures that a climate-cooling effect (negative RF_bio_) yields a positive ME, aligning higher ME values with greater mitigation benefit. This makes ME a consistent and intuitive indicator of strategy performance across time horizons.

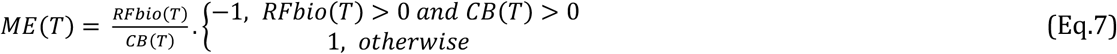

ME is calculated across different time horizons (85, 185, and 285 years), providing a metric that relates the remaining or effectively removed biogenic emissions from the atmosphere to the total carbon balance. The higher the ME value, the greater the climate benefit delivered per unit of carbon captured, making it a useful comparative indicator of mitigation strategy performance.

#### 2.5.1 Statistical Analysis of Mitigation Potential

The effects of forest management, climate scenarios, wood use, and substitution factors were statistically evaluated across a range of carbon-related response variables in the forest sector. Carbon ecosystem fluxes (i.e., GPP, R_ECO_, and NEE) were analyzed with respect to forest management and climate, while wood emissions, substitution effects, wood balance, and FSB were evaluated in relation to all four factors.

We used linear mixed-effects models (LMEs) to regress GPP, RECO, NEE, and FSB against five key drivers: Management, Wood Use (WU), Substitution (SUB), Climate, and Century. These models were used to assess the influence of each factor and their interactions on ecosystem fluxes and forest sector balances. For wood emissions, substitution effects, and wood balance, we applied generalized additive mixed models (GAMMs) to account for potential nonlinear relationships. Model selection was guided by residual diagnostics, including assessments of linearity, normality, and homoscedasticity. The factors and their acronyms, used consistently throughout the manuscript, are summarized in **Table 1**. Full model specifications, transformations, and diagnostic results are provided in **S2**.

**Table 1:** Overview of the factorial design used in the simulation experiment, showing the five key factors and their respective levels along with their corresponding modality acronyms.

| Factor | Levels | Modality acronyms |
| --- | --- | --- |
| Management | 5 | TM, WOOD, BIOE, ADAPT, TRANS |
| Wood use | 4 | BAU, RR20, LS20, RRLS20 |
| Substitution | 5 | SUB0, SUB25, SUB50, SUB75, SUB100 |
| Climate change scenario | 3 | CUR, SSP1-2.6, SSP5-8.5 |
| Century | 3 | C1 (2015-2099), C2 (2100-2199), C3 (2200-2300),<br>ALL (2015-2300) |

#### 2.5.2. Consistency between carbon and climate mitigation rankings and the concept of mitigation efficiency

To evaluate the relationship between CB and RF_bio_ outcomes, a linear regression analysis was conducted, modeling RF_bio_ as a function of CB across all studied scenarios. Model fit was assessed using the coefficient of determination (R^2^) and statistical significance (p-value). ME was calculated for each combination of climate scenario, management scheme, substitution pathway, and time horizon (H1: 2015–2100; H2: 2015–2200; H3: 2015–2300). To assess the sensitivity of ME to scenario context and temporal scope, we conducted a descriptive statistical comparison across each factor. Patterns in ME and CB were summarized using means, with scenario-specific anomalies (e.g., negative ME values) noted.

Additionally, an in-depth analysis of two forest mitigation options that exhibited the two most divergent ME was conducted. This comparison focused on long-term FSB dynamics, RF_bio_ patterns, time in a net emissions regime, and disturbance-recovery cycles, to elucidate the underlying trade-offs between cumulative carbon benefits and the timing and distribution of emissions.

## 3. Results

### 3.1. Forest sector balance (FSB)

FSB, defined as the sum of Net Ecosystem Exchange (NEE) and the wood-product balance (Wbalance; i.e., substitution effects minus emissions from wood decay in disposal sites), varied significantly with climate scenarios, management strategies, and across centuries. Climate effects on FSB were not consistent across forest management types, and both drivers showed strong century-dependent patterns (**see Table S2.4).** Although the displacement factors (DF) had a low overall effect size (η² = 0.03, p<0.001), its influence on FSB was substantial in specific scenarios, particularly those representing rapid decarbonization pathways. Over the full simulation period, SSP5-8.5 yielded the lowest FSB (0.07 ± 0.01 tC ha^-1^ year^-1^), while SSP1-2.6 and current climate (CUR) resulted in significantly higher values (0.75 ± 0.01 and 0.64 ± 0.01, respectively). The ranking of scenarios changed over time: SSP5-8.5 initially outperformed others but declined in the 22nd and 23rd centuries, ultimately ranking lowest (**see Figure 6**). By contrast, SSP1-2.6 and CUR remained more stable across centuries with lower inter-temporal variability.

**Figure 6:**
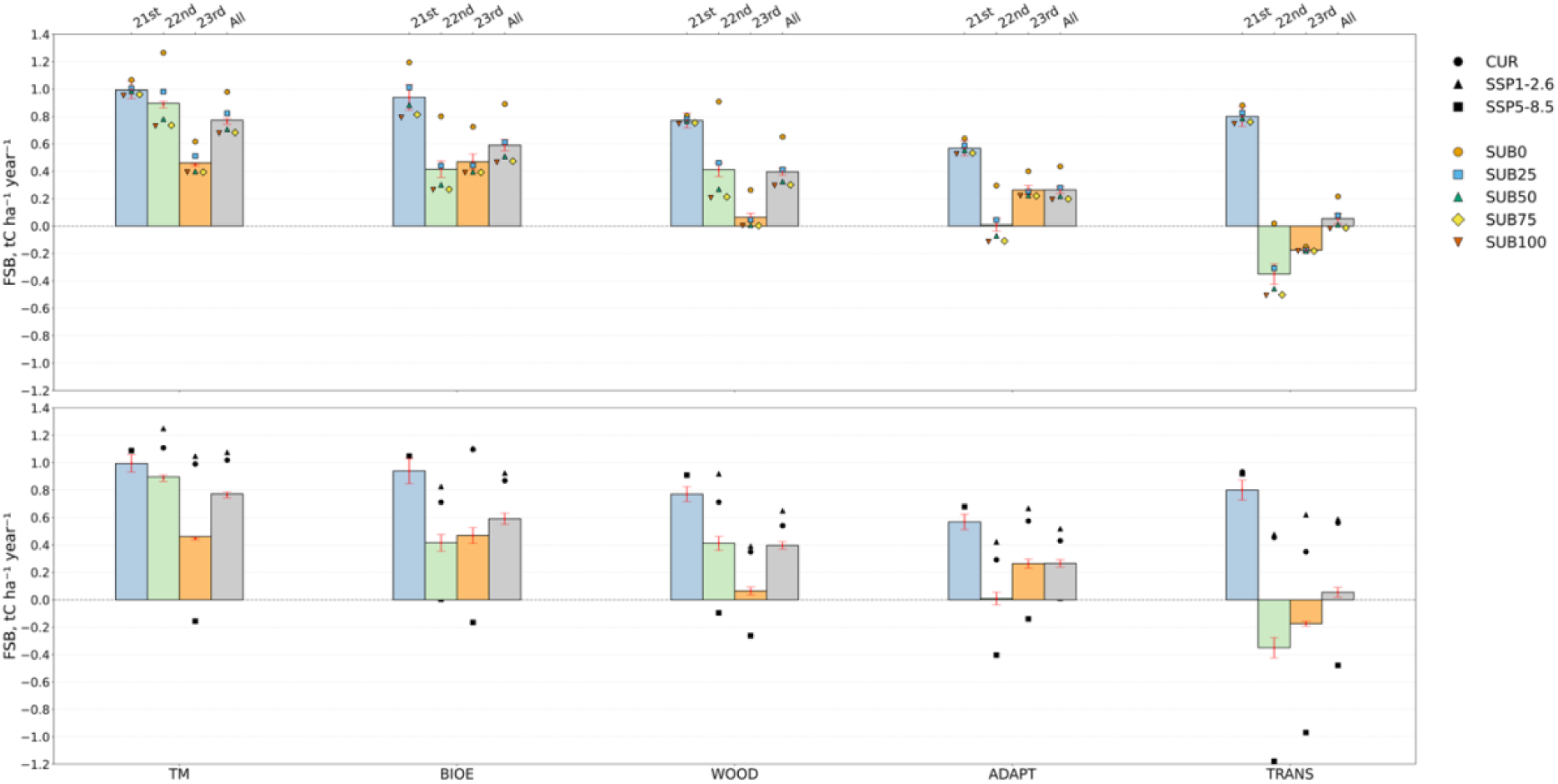
Mean FSB, in tC ha^-1^ year^-1^ for five forest management strategies (TM, ADAPT, BIOE, TRANS, and WOOD), averaged across two climate scenarios (SSP1-2.6 and SSP5-8.5), shown for the 21st, 22nd, and 23rd centuries, as well as for the full period combined. Red error bars indicate the 95% confidence interval of FSB across climate scenarios. The upper panel shows FSB values by substitution scenarios, while the lower panel shows FSB values by climate scenario.

Management strategies showed clear performance gradients mainly driven by NEE. Modular harvesting (TM) and bioenergy (BIOE) produced the highest forest sector balance (FSB; 0.76 ± 0.02 and 0.59 ± 0.04), supported by relatively strong sinks (NEE: –1.25 ± 0.05 and – 1.10 ± 0.08). Promotion of long-lived wood products (WOOD) and forest adaptation management (ADAPT) showed intermediate performance (FSB: 0.39 ± 0.02 and 0.26 ± 0.02) associated with weaker sinks (NEE: –0.78 ± 0.05 and –0.54 ± 0.05). In contrast, transition to passive rewilding (TRANS) consistently produced the lowest FSB (0.05 ± 0.03), declining to negative values under SSP5-8.5 (–0.47 ± 0.03) due to forest ecosystem turning into carbon source (NEE: +0.83 ± 0.01) (**see Figure 6**). Decreases were most severe in TRANS and ADAPT, where net carbon source behavior emerged in the third century.

DF, which modulates substitution levels over time, had a major influence on Wbalance and thus on FSB. Under the baseline pathway (SUB0), the substitution effect was maintained throughout all centuries, while decarbonization pathways (SUB25–SUB100) showed a progressive decline in substitution, reaching zero by 2155 (SUB25), 2085 (SUB50), 2062 (SUB75), and 2050 (SUB100), respectively. These trajectories form three distinct temporal patterns: persistent (SUB0), medium-lived (SUB25), and short-lived (SUB50–SUB100), which resulted in markedly different Wbalance outcomes.

Component analysis indicated that NEE accounted for ∼53–65% of FSB, particularly high under TM. Substitution effect was lower in terms of contribution to the FSB but variable, peaking under the SUB0 displacement pathway (18%), decreasing under SUB25 (7%), and falling below 3% for SUB50–SUB100, which caused a (0.28 ± 0.07 tC ha^-1^ year^-1^; 53 ± 25%) reduction in FSB compared to SUB0 (**see Table S2.5**).

### 3.2. Radiative forcing & mitigation efficiency (RF_bio_ and ME)

Forest management scenarios with a more negative carbon balance (CB) also tended to yield the greatest climate benefits. A linear regression, performed using outcomes from all studied scenarios, confirmed that CB is a strong predictor of the radiative forcing from biogenic emissions (RF_bio_; R² = 0.95; *p* < 2.2e–16; see **Table S2.9**). Thus, the results and analyses previously conducted on CB are also applicable to RF_bio_. However, this relationship is not always strictly linear. In some exceptional cases where CB values are close, a mitigation option with a slightly less negative CB produced a more favorable RF_bio_. This was observed, for instance, under the SUB25 decarbonization pathway at horizon H2 (2015-2200): the TRANS scenario had a slightly higher CB (–128 ± 0.91 tC ha^-1^) than WOOD (–122 ± 2.75 tC ha^-1^), yet WOOD achieved a more favorable RF_bio_ (–66 ± 1.33 vs. –60 ± 0.13 tC ha^-1^) and a higher ME (0.54 ± 0.00 vs. 0.47 ± 0.002). At horizon H3 (2015-2399), this contrast became even more pronounced: although TRANS reached a more negative CB (–162 ± 0.92 tC/ha) compared to WOOD (–155 ± 2.05 tC ha^-1^), it resulted in a substantially less favorable RF_bio_ (–66 ± 0.70 vs. –73 ± 0.20 tC ha^-1^) and a lower mitigation efficiency (ME; 0.41 ± 0.002 vs. 0.57 ± 0.002). ME showed strong sensitivity to both climate conditions and forest management practices.

Under CUR and SSP1-2.6, ME remained relatively stable across management schemes, typically ranging from 0.4 to 0.7 (**Figure 7** and **Table S2.10**). In contrast, under the SSP5-8.5 climate scenario, the ME of different management scenarios was more variable and often with lower efficiencies, with the efficiency of several scenarios falling below zero, particularly ADAPT and TRANS under H2 and H3 (**Figure 7** and **Table S2.10**), indicating that these scenarios were contributing to climate warming rather than cooling it. The time horizon (H1, H2, and H3) also influenced ME: across all climate scenarios, the short-term horizon (H1) consistently resulted in higher efficiency values (>0.6) compared to the longer-term time horizon (H2 and H3).

**Figure 7:**
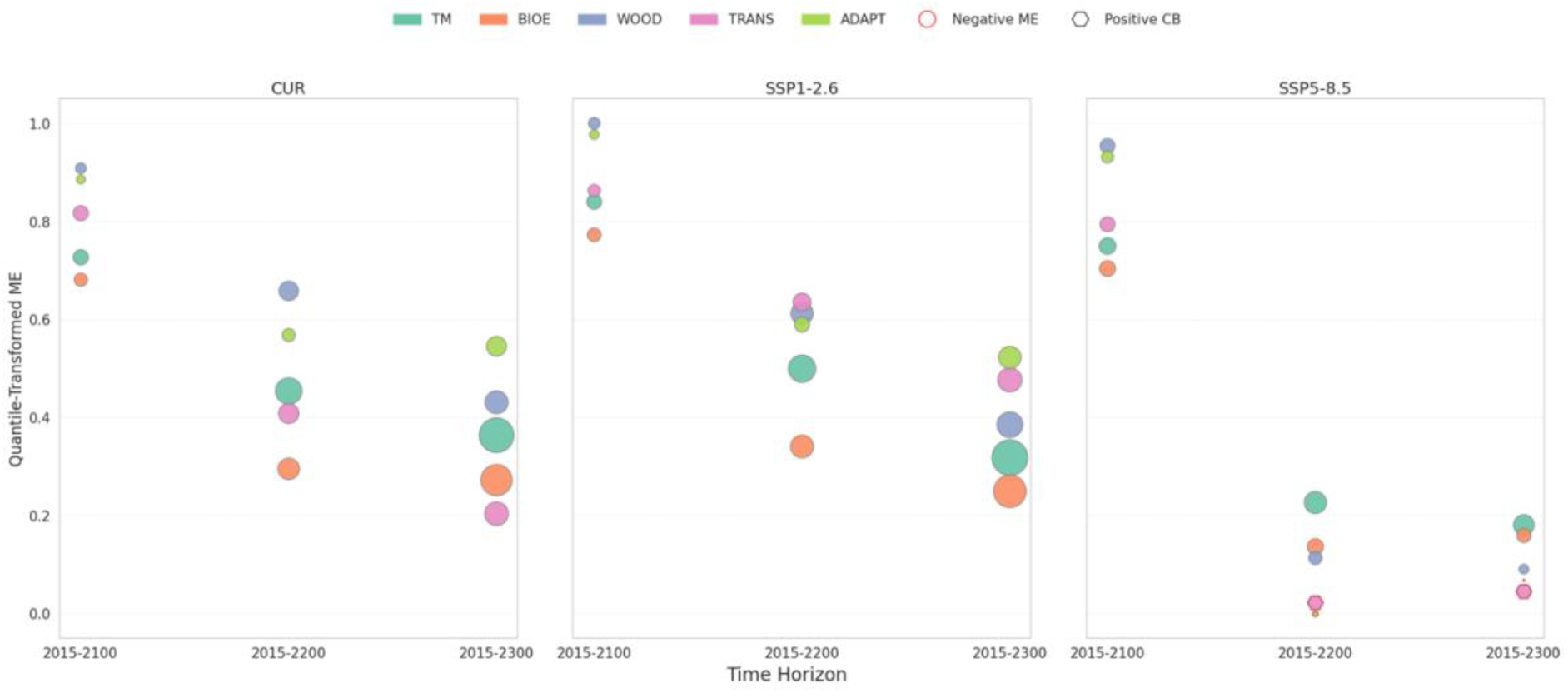
CB and ME under various forest management and climate scenarios. Circle size scales with CB via −0.9 × *CB*^1.3^; ME is quantile-transformed, both for improved visibility and comparability across scenarios. Red borders indicate negative ME; hexagons show positive CB (carbon source sector). Subplots represent climate scenarios, with circles colored by management schemes.

Interestingly, a more negative CB did not always equate to high efficiency. In CUR and SSP1-2.6, forest projects with stronger (more negative) CB often showed lower or slightly lower efficiency, whereas the opposite was true under SSP5-8.5, where stronger CB tended to support higher ME. For example, ADAPT (0.53, SSP1-2.6, H2) and WOOD (0.54, SSP1-2.6, H2) sometimes outperformed TM (0.49, SSP1-2.6, H2) or BIOE (0.45, SSP1-2.6, H2) in efficiency despite having less negative CB values.

These dynamics were further supported and extended by generalized additive models (GAMs; **Figures S2.1-S2.2**), which demonstrated that, in addition to CB, both the magnitude and duration of emissions play dominant roles in determining ME. Under the SSP5-8.5 scenario, emissions magnitude and CB were strongly positively correlated (r = 0.85), indicating that scenarios with more negative CB generally exhibited low emissions. In contrast, a weak negative correlation (r = –0.28) was observed under CUR and SSP1-2.6, suggesting no clear relationship between CB and emissions magnitude. In the ADAPT and TRANS scenarios, under the SSP5-8.5 climate scenario, positive CB values resulted in sustained net emissions and positive contributions to radiative forcing.

### 3.3. In depth-insights from BIOE and WOOD strategies

Across the simulation period (2015–2300), the BIOE management scenario consistently achieved a more negative CB than WOOD. By 2300, BIOE reached –337 tC ha^-1^, while WOOD reached –206 tC ha^-1^. ME, however, remained higher in WOOD throughout. Both scenarios began with comparable ME in 2100 (0.62 for BIOE, 0.63 for WOOD). By 2200, WOOD retained a ME of 0.54, while BIOE declined to 0.46. In 2300, ME values were 0.47 for WOOD and 0.44 for BIOE (see **Figure 8**).

**Figure 8:**
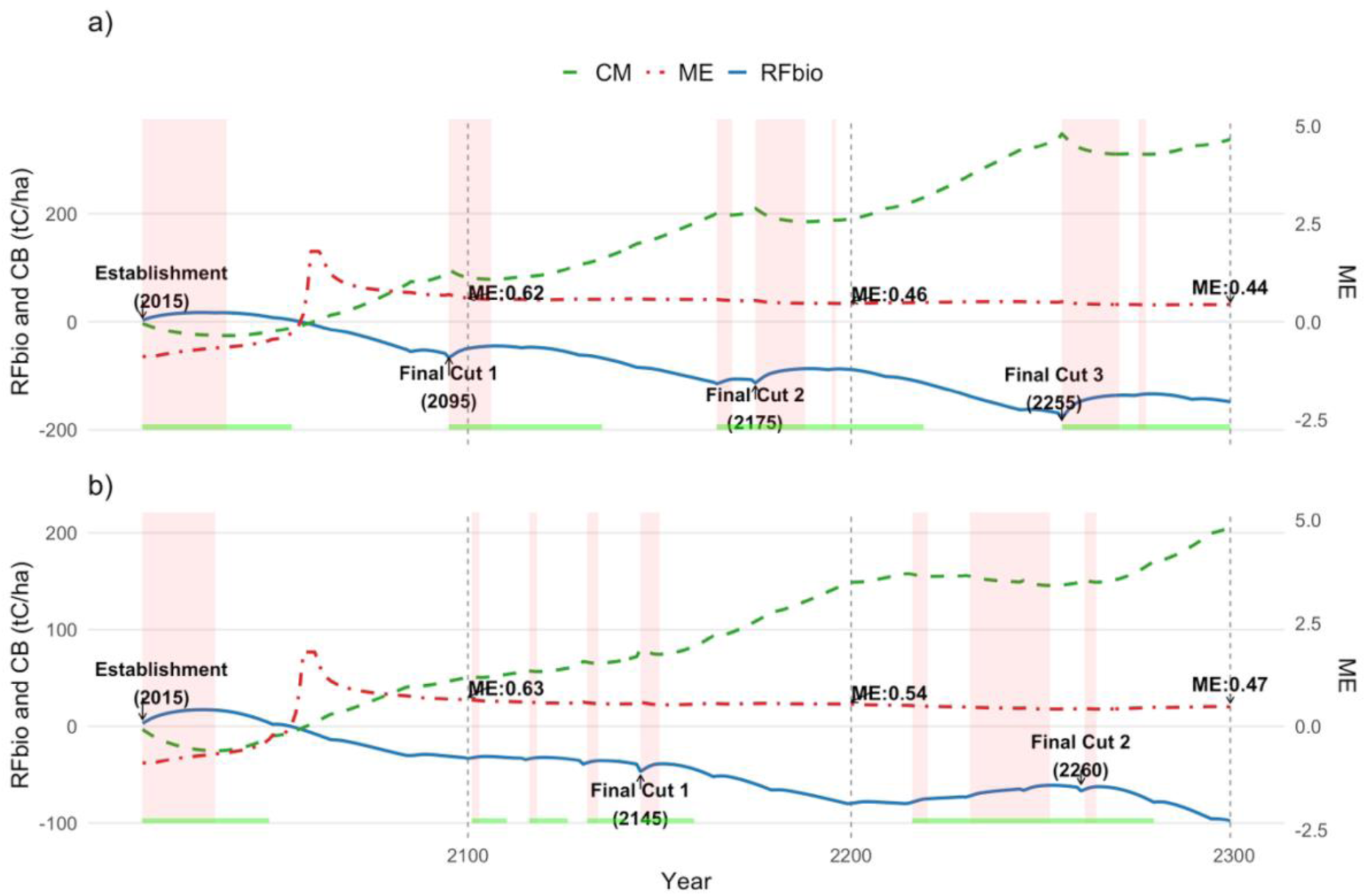
Temporal dynamics of RF_bio_, CB, and ME under BIOE (panel a) and WOOD (panel b) management scenarios from 2015 to 2300. Shaded red bands indicate periods when the forest sector is a net source (FSB<0). Green horizontal bands denote the number of years required for RF_bio_ to return to its pre-harvest level.

BIOE followed an 80-year shelterwood rotation, resulting in three major final harvests occurring in 2095, 2175, and 2265. WOOD used a 130-year rotation, with final cuts in 2145 and 2260. The first BIOE final harvest in 2095 led to emissions of approximately 17.3 tC ha^-1^, with RF_bio_ returning to pre-disturbance levels by 2135, indicating a 40-year payback period. For WOOD, the first final cut in 2145 resulted in emissions of 5.6 tC ha^-1^, with RF_bio_ returning to net cooling within 15 years. Following these major harvests, NEE in WOOD returned to negative values (net sink) within 4 years, whereas BIOE remained a net carbon source (NEE > 0) for approximately 10 years.

In the three years immediately following each final cut, RF_bio_ increased sharply in BIOE, while the increase in WOOD was more moderate. This early post-harvest rise in RF_bio_ coincided with higher volumes of short-lived HWPs (i.e., firewood and paper) in BIOE; 37.25 m³ ha^-1^, 38.51 m³ ha^-1^, and 37.88 m³ ha^-1^ after its respective final harvests, compared to 23.70 m³ ha^-1^ and 14.82 m³ ha^-1^ in WOOD.

Emission timing and magnitude differed between the two scenarios. In BIOE, emissions were concentrated in a limited number of high-intensity events, associated with initial establishment and final harvests. These events corresponded to periods where GPP was low and ecosystem respiration (R_ECO_) remained high although the carbon outflow from the forest ecosystem (Rh_onsite_) decreased proportionally to GPP. Additional immediate emissions from firewood combustion or from products at their end-life incineration (i.e., paper) were also observed during these intervals. In WOOD, emissions were more evenly distributed and of lower magnitude, arising from thinning, regeneration phases, and smaller final cuts. RF_bio_ patterns in WOOD were more stable, with shorter durations of net positive forcing following harvest events.

## 4. Discussions

Our results suggest that effective forest-sector climate mitigation requires more than maximizing carbon balances. It demands a systems perspective integrating forest dynamics, wood use, and evolving socio-technical conditions. Over a multi-century assessment, we found that while the forest sector balance (FSB) is strongly influenced by climate and management, mitigation efficiency (ME) depends more on emission timing and scale. Substitution effects, particularly the contrast between the baseline pathway (SUB0) and the degressive pathways (SUB25-SUB100) proved substantial, while HWP lifespan and recycling rate had negligible long-term impact. Notably, scenarios with the lowest carbon balance (CB) did not always deliver the greatest climate efficiency. ME declined over time, emphasizing the need to consider temporal dynamics. All management options eventually showed declining carbon sink capacity under SSP5-8.5, with TRANS, ADAPT, and WOOD becoming net sources.

### 4.1. Management strategies, wood use, and substitution pathways

Forest management has a strong influence on FSB. Intensive silvicultural practices, such as TM and BIOE, consistently maintained high FSB across climate projections. (Giasson et al., 2023) showed that harvest scenarios with high intervention and regular regrowth cycles sustain stronger carbon sink than low-harvest or unmanaged systems. Furthermore, our results showed that shifting from active to passive management under the TRANS scenario significantly reduced FSB in the long term, especially under contrasted climate conditions where forest carbon dynamics approached equilibrium. These findings align with (Pukkala, 2017), who reported that unmanaged forests exhibit a long-term decline, nearing zero around 200 years post-establishment. Beyond carbon storage, management also influences wood volume and characteristics, affecting substitution potential.

Recent studies increasingly challenge static DFs, advocating for dynamic approaches that reflect evolving technological, industrial, and policy contexts (I. Boukhris et al., 2025; Brunet-Navarro et al., 2021; Hurmekoski et al., 2023). Our dynamic modeling confirms these concerns. In scenarios where DFs reaches zero by 2050, substitution benefits disappear entirely, resulting in a 53% lower FSB by 2300 compared to the SUB0 trajectory. This mirrors projections by Brunet-Navarro et al. (2021), who estimated substitution benefits may decrease by approximately 30% by 2030 and up to 96% by 2100, in line with the decarbonization goals of the Paris Agreement.

Conversely, wood-use parameters like product lifespan and recycling rate had no discernable long-term impact on FSB. Over 285 years, stored carbon in HWPs is eventually emitted through decay, regardless of whether turnover occurs at 40 or 60 years. Although extended lifespans and higher recycling rates may reduce emissions in the short- to medium-term, as shown by (Brunet-Navarro et al., 2016), who projected up to 5 Mt CO_2_ yr^-1^ savings in the EU-28 with lifespan increases of 19.54% or a 20.92% recycling boost by 2030, these effects diminish across multi-century timescales. Such results highlight the need to align interventions with the analytical timeframe. Short-term gains may appear significant but are absorbed over longer horizons when delayed emissions fall within the evaluation period.

### 4.2. Carbon sink instability under high-emission scenarios

The shift to net-carbon neutrality or a carbon source state by the last considered century under SSP5-8.5 marks a critical loss of ecosystem resilience, driven by prolonged structural degradation and physiological stress from high temperatures. Although precipitation increases during this period, and vapor pressure deficit stays moderate (∼0.825 ± 0.1 kPa), GPP declines sharply across all strategies. This drop is more likely due to heat-induced impacts on photosynthetic processes, such as enzyme deactivation, increased photorespiration, and reduced carboxylation efficiency (Oliver et al., 2022; Qu Y et al., 2023). As biomass and metabolic activity decline, both Ra and Rh fall due to limited substrate and structural loss, but GPP declines faster, leading to sustained positive NEE in the final simulation century. This aligns with Earth System Model outputs under SSP5-8.5 which show that warming suppresses photosynthesis as temperatures exceed physiological thresholds, particularly in temperate and boreal forests. The summer GPP-temperature correlation weakens over the 21st century, turning significantly negative by 2070 (Chen and Peng, 2025; Zhang et al., 2022) also found that 33% to 63% of global vegetated areas show declining productivity sensitivity to temperature, with the most severe declines under SSP5-8.5.

### 4.3. Climate-Carbon dynamics and mitigation efficiency

The strong correlation observed between carbon balance (CB) and radiative forcing from biogenic emissions (RF_bio_) reflects the key role of long-lived CO_2_ removals in determining climate impacts of forest-based mitigation. This aligns with carbon cycle theory and is consistent with impulse response functions (IRFs) such as the Bern model (Joos et al., 2013), where most climate benefit arises from sustained atmospheric CO_2_ reduction (Archer et al., 2009; Masson-Delmotte et al., 2021). However, the CB-RF_bio_ relationship is not strictly proportional. Our results show that under the SUB25 decarbonization pathway at both H2 and H3, the WOOD scenario achieved a lower (better) RF_bio_ than TRANS, despite similar or less negative CB. This suggests CB alone is insufficient basis for assessing climate performance, and other factors also influence radiative forcing.

A key reason for this explanation for the discrepancy is the timing of carbon fluxes. CB summarizes total emissions and removals but masks their temporal distribution. Many scenarios involve non-monotonic dynamics, with initial emissions from establishment, thinning, or harvest followed by delayed regrowth. Early emissions exert greater radiative forcing due to the CO_2_’s long atmospheric lifetime cumulative warming (Allen et al., 2010; Matthews and Caldeira, 2008). Although IRF is linear for pulses, time-distributed fluxes interact non-linearly with the climate system. RF_bio_, which captures these dynamics, more accurately reflects climate impact than CB alone (Sierra et al., 2021; Tanaka and O’Neill, 2018).

Nevertheless, neither CB nor RFbio capture the carbon “cost” of achieving mitigation. We therefore introduced the mitigation efficiency (ME; RFbio divided by CB). ME indicates how effectively carbon removal translates into avoided radiative forcing. For instance, BIOE had a more negative CB (−339 tC ha^-1^) than ADAPT (−162 tC ha^-1^), but a lower ME (0.44 vs. 0.49), meaning less efficient climate benefit. BIOE also required 17 silvicultural interventions and 3 final cuts over 300 years, compared to 14 interventions and 2 final cuts in ADAPT. Thus, more intensive management does not always yield proportionally greater mitigation and may entail higher ecological or operational costs.

ME declined consistently as the time horizon extended. Initially, forests sequester carbon in biomass and soils, but later interventions transfer carbon to HWPs, which eventually decay and emit CO_2_. Over time, emissions from earlier cycles accumulate, reducing net benefit. This dynamic, “sequester now, emit later” (Guest et al., 2013), underscores the value of ME as a complementary metric that integrates both climate effectiveness and implementation sustainability.

### 4.4. Uncertainty, Model assumptions, and Long-Term Implications

Uncertainties within the modeling framework stem from initialization, process representation, and parameterization ((Hamish) Kimmins et al., 2008; Reyer et al., 2020). The 3D-CMCC-FEM forest growth model, while effective for stand-level analyses, simplifies spatial heterogeneity and omits landscape-scale dynamics, potentially underrepresenting key interactions such as seed dispersal and spatially explicit disturbance spread (Patacca et al., 2023; Seidl et al., 2011). It also omits processes like tree establishment, disturbance-induced mortality, and species composition are not represented, limiting its ability to capture ecological feedback under climate change. Adaptive responses, including species acclimation, migration, and phenological change, are excluded or overly simplified, despite their growing relevance over multi-century timescales (Issam Boukhris et al., 2025; Nicotra et al., 2010).

On the HWPs side, the TimberTracer model uses fixed bucking rules and constant product allocations, without accounting for future technological shifts, circular economy developments, or changing consumer demand (Harju, 2022; Olšiaková et al., 2016). This restricts its ability to reflect how innovation or policy changes could influence product lifespans, recycling rates, or end-of-life treatment.

Additional uncertainty comes from external drivers. Climate projections diverge significantly after 2100 (Hausfather and Peters, 2020), and future land-use and forest management pathways depend on socio-economic and policy trends that are inherently hard to predict (Popp et al., 2017). DF estimates used to quantify substitution benefits are also highly uncertain and context-specific (Leskinen et al., 2018; Pingoud et al., 2010). Their magnitude will likely evolve as energy systems decarbonize and industry shifts to lower-carbon alternatives (Gustavsson et al., 2017).

Although the Vallombrosa site represents a productive temperate European forest, this study aims to identify general mechanisms rather than universal prescriptions. The relative ranking of management strategies observed here should be interpreted as site-dependent. In highly productive temperate systems, rapid regrowth can offset harvest emissions, whereas in boreal forests slower recovery may extend emissions payback times and favor lower harvest intensities. Conversely, in drought-prone Mediterranean regions or disturbance-dominated mountain forests, mortality dynamics may outweigh planned interventions. Nevertheless, the structural relationships identified in this study appear robust: carbon balance alone does not determine climate benefit, mitigation efficiency depends on emission timing and magnitude, and declining substitution factors systematically reduce long-term climate mitigation. These mechanisms arise from carbon-cycle dynamics rather than site-specific properties and are therefore expected to apply broadly across forest contexts. Considering this, EU-level mitigation assessments based solely on carbon stock indicators may overlook important climate-impact differences among strategies. Rather than prescribing uniform management targets, policy frameworks may benefit from approaches that evaluate mitigation performance using temporally explicit metrics and that allow regional adaptation to ecological conditions and evolving substitution effectiveness.

## Conclusions

This study critically examined the long-term climate mitigation potential of the forest sector by integrating forest growth, wood-use, and substitution dynamics within a unified modeling framework. Our aim was not only to assess carbon balances but to evaluate whether commonly used indicators accurately reflect true climate benefit over multi-century timescales.

Using a coupled modeling approach applied on a *P. nigra* forest in Italy, we simulated a broad range of futures across five forest management strategies, four wood-use patterns, and five substitution decay scenarios under three contrasting climate pathways. While CB and RF_bio_ were generally aligned, they captured different aspects of climate impact. CB reflects net carbon flows, but not their atmospheric effect; RF_bio_ represents the warming or cooling influence overtime. To bridge this gap, we introduced ME, a metric that relates carbon flows to actual contribution to actual mitigation effectiveness.

Our findings show that scenarios with the highest CB (e.g., BIOE and TM) do not always deliver the greatest climate benefit, especially when emissions are front-loaded or persistent. Meanwhile, strategies like WOOD and ADAPT, which extended carbon residence times in both forest and technosphere, achieved higher ME under SSP1-2.6. However, under the SSP5-8.5 scenario, even these strategies became net radiative sources after 2200 due to declining sink capacity.

Substitution benefits, often assumed constant, declined significantly under degressive assumptions, reducing the overall forest sector mitigation by up to 53%, especially in high harvest volumes. While substitution played a secondary role compared to climate and management, its sensitivity to decarbonization pathways highlights the risk of overreliance on this lever.

Taken together, these findings highlight that effective forest-sector mitigation requires more than achieving high carbon balances. It also depends on minimizing the magnitude, timing, and persistence of emissions, and on reevaluating the assumptions behind substitution effects. Metrics like RF_bio_ and ME provide valuable tools to capture these temporal and systemic dynamics, offering a more nuanced, climate-relevant basis for evaluating forest-sector strategies.

Ultimately, meaningful forest-sector contributions to climate neutrality require assessment not just of carbon flows, but of their actual atmospheric impact over time.

## Appendix

**Figure A.1:**
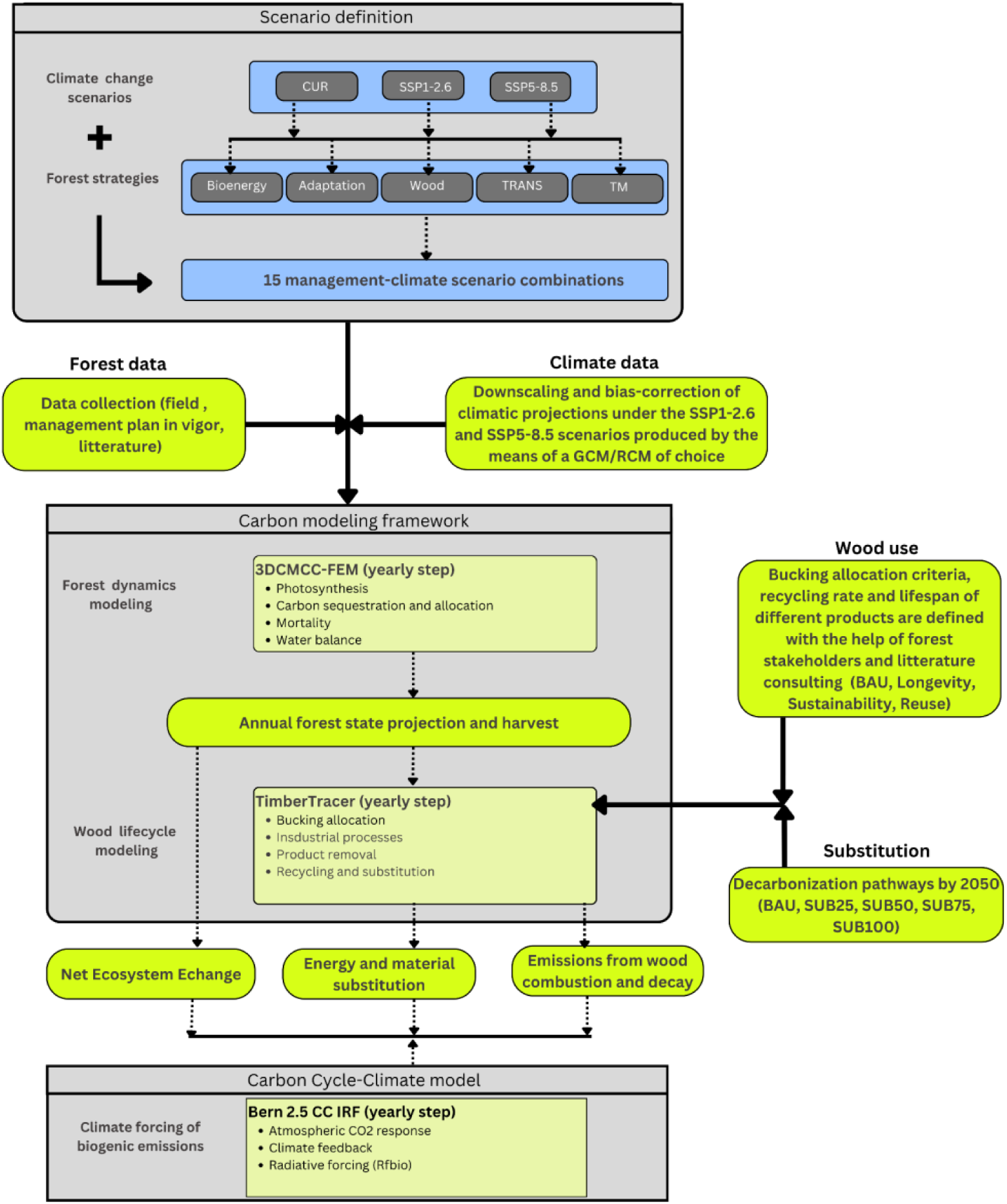
Integrated modeling framework used to evaluate the long-term climate mitigation potential of a managed *Pinus nigra* forest (2015-2300). The framework combines forest dynamics (3D-CMCC-FEM), wood lifecycle modeling (TimberTracer), and an impulse response function (Bern 2.5 CC IRF). Simulations cover multiple scenario combinations (for management, climate, wood-use, and substitution). Outputs include temporal net ecosystem exchange, substitution effect, and emissions from wood combustion and decay, which are used to compute forest sector balance and radiative forcing from biogenic CO_2_ emissions.

## Credit authorship contribution statement

**Issam Boukhris:** Writing - original draft, Writing - review & editing, Methodology, Visualization, Validation, Investigation, Data curation, Conceptualization, Project administration. **Francesco Cherubini:** Writing - review & editing, Methodology, Resources, Formal analysis, Resources. **Alessio Collalti:** Writing - review & editing, Validation, Software, Formal analysis, Resources. **Daniela Dalmonech:** Validation, Software, Resources. **Christian Vonderach:** Writing - review & editing, Investigation, Validation, Formal analysis. **Francesca Giannetti**: Writing - review & editing, Resources, Formal analysis. **Gina Marano:** Writing - review & editing, Methodology, Formal analysis **Said Lahssini:** Writing - review & editing, Formal analysis, Software. **Monia Santini:** Writing - review & editing, Software, Funding acquisition. **Riccardo Valentini:** Formal analysis, Supervision, Funding acquisition.

## Data availability Statement

Data are available from the corresponding author upon reasonable request.

### Acknowledgment

Issam Boukhris would like to thank the University of Tuscia and the CMCC Foundation, both based in Italy, for supporting his Ph.D. thesis, of which this work is a part. He also extends his gratitude to Professor Gherardo Chirici and Dr. Elia Vangi from the University of Florence, Vincenzo Saponaro from the CNR of Italy, Colonel Giuliano Savelli and Maréchal Giovanni Galipò from the local forest services of Vallombrosa for their technical and logistical support. Special thanks are also due to Dr. Khalil Misbah from the IRD of Toulouse, as well as to Adnan Yousaf and Renato Zompanti from the University of Tuscia, for their invaluable assistance during fieldwork.

