## Supplementary Material 1 for "Long-term forest-sector mitigation and radiative forcing under contrasting management, climate, and substitution pathways"

**Supplementary 1: Model setup and parametrization**

**1. Models description**

1.1. **3D-CMCC-FEM**

The 3D-CMCC-FEM (‘*Three Dimensional – Coupled Model Carbon Cycle – Forest Ecosystem Model*’, v.5.7) is a process-based model integrating biogeochemical, biophysical, and physiological processes to predict carbon, nitrogen, energy, and water fluxes, as well as stand development dynamics that influence changes in forest ecosystem fluxes and stocks (Collalti et al., 2014; Dalmonech et al., 2024; Testolin et al., 2023; Vangi et al., 2024). It is designed to simulate physiological and hydrological processes with species-specific resolution on daily, monthly, and annual scales, depending on the simulated process. The 3D-CMCC-FEM model requires a set of input data for its initialization which includes: (1) species-level model parameters (species-specific eco-physiological and allometric characteristics that can be derived from forest inventories and literature), (2) site data (e.g., latitude, altitude, soil properties), (3) initial stand structural data (e.g., DBH, tree height, stand density, age), (4) climatic data (meteorological data at a daily time step, e.g., mean daily incident solar radiation, mean daily air temperature). To ensure equilibrium in soil carbon pools before running simulations, a spin-up process was conducted by cycling climate data from 1991-2020 over 2000 years.

Parameterization for *Pinus nigra* was based on detailed field data collected from permanent sample plots. Where species-specific physiological parameters were unavailable, proxies were drawn from ecologically similar species (*P. pinaster*, *P. sylvestris*, *P. ponderosa*) and other evergreen conifers. The complete list of parameters and references is provided in Supplementary Table S2. Daily climate data covering the simulation period (2005–2023) were obtained from the regional hydrological service of the Tuscany Region ([https://www.sir.toscana.it](https://www.sir.toscana.it/)). These were downscaled to plot level using the MT-CLIM microclimate model (Thornton and Running, 1999), with adjustments for local topographic variation. Model validation was conducted using both structural and flux-based metrics. Structural outputs, such as DBH, tree height, and stand density, were compared against annual field measurements. Monthly flux outputs, specifically GPP and NPP were validated using satellite-derived estimates from the MODIS MOD17A2HGF dataset (Running & Zhao, 2021).

**1.2. TimberTracer**

The TimberTracer (TT; Boukhris et al., 2025) is a wood product model (WPM) based on the material flow method which comprehensively accounts for and simulates the temporal dynamics of GHG emissions and removal across all carbon pools outside the forest, including changes in HWPs and disposal sites. Furthermore, the model considers the substitution effect and captures the temporal dynamics of material and energy substitutions. TT model requires a set of inputs and parameters necessary for its initialization: (1) the bucking allocation criteria are standardized rules used to sort solid-wood products by quality for different end uses, and they can be applied regardless of the log source or sawmill (Josza et al., 1994). Potential wood products from black pine stems were inventoried by consulting five sawmill industry experts, and the compilation of all possible products was retained for this study. Furthermore, the bucking criteria used in this study are those commonly found in the literature (CTBA, 2001) (Please refer to Table 2); (2) the transformation efficiency of each log category, a geometric yield as well as the loss reallocation were defined with the assistance of sawmill industry professionals (Please refer to [Tables](https://www.biorxiv.org/content/10.1101/2024.01.24.576985v2.full#T2) 3 and 4); (3) for the BAU recycling rate we suggest that the recycling rate of waste wood products was constant at 10% during the time horizon while lifespan of each product was reviewed from published studies (Burschel et al., 1993; E. Skog & A. Nicholson, 1998; Eggers, 2002; Karjalainen et al., 1994; Masera et al., 2003; Nabuurs, 1996; Pingoud et al., 2003) (Please refer to [Tables](https://www.biorxiv.org/content/10.1101/2024.01.24.576985v2.full#T2) 4 and 5); (4) displacement factors for the substitution were derived from the literature (Sathre & O’Connor, 2010; Suter et al., 2017) Please refer to [Table](https://www.biorxiv.org/content/10.1101/2024.01.24.576985v2.full#T5) 6); (5) TT model dendrometric parameters, including stand structure, crown base height equation, diameter-to-height parameters, were fitted to the forest data collected from the NFI plots of the Vallombrosa forest, the taper model from (Boukhris et al., 2024), while the species wood density was derived from literature (Guner & Çömez, 2017).

2. **Model validation**

To parameterize and validate the forest model, we used data from 14 permanent sample plots that provided detailed ground-based measurements of tree structure and growth (dendrometric observations). In addition to these field data, we incorporated satellite-based estimates of forest productivity to evaluate the model's performance. Specifically, we used gross primary productivity (GPP) and net primary productivity (NPP) data from the MODIS MOD17A2HGF version 6 product, which provides 8-day composite values at a spatial resolution of 500 meters. This remote sensing dataset allowed us to assess how well the model reproduced key ecosystem carbon fluxes over time. To ensure the reliability of the satellite data, we filtered it to exclude pixels affected by cloud cover, poor sensor conditions, or low confidence, using the quality control indicators included in the MODIS product. Additionally, because our sample plots (530 m^2^) are much smaller than a MODIS pixel (500 × 500 m), we applied further spatial filtering to ensure land cover consistency. Specifically, we retained only those pixels in which non-forest land cover was less than 5%, and in which coniferous forest made up more than 75% of the forested area. This step was essential to ensure that the selected MODIS pixels accurately reflected the forest conditions of interest. The filtered satellite data were then aggregated to monthly and annual time scales to match the model’s temporal resolution and enable meaningful comparison between observed and simulated productivity**.**

The model was parameterized to simulate the growth and development of black pine stands, using field data where available to calibrate allometric parameters, and supplementing with the broad published literature on the species (Debain et al., 2007; Fernández-Martínez et al., 2017; Isajev et al., 2004; Nagel et al., 2023; Navarro-Cerrillo et al., 2016). For physiological parameters lacking specific data for *Pinus nigra*, proxy data from ecologically-similar species were used in the following order of preference: *Pinus pinaster* (Nunes et al., 2015), *Pinus sylvestris* (ALBREKTSON, 1984; Chmura et al., 2013; Kuusinen et al., 2014; Mencuccini & Grace, 1995; Tatarinov & Cienciala, 2006; van Hees & Bartelink, 1993; Vanninen et al., 1996), *Pinus ponderosa* (Mitra et al., 2020), or more broadly, evergreen species (Arora & Boer, 2005). Detailed parameters and their respective sources are provided in Table 7.

The 3D-CMCC-FEM model requires daily climatic inputs such as solar radiation (MJ m^-2^), temperature (°C), precipitation (mm), and vapor pressure deficit (hPa). Data spanning the period from 2005 to 2023 were sourced from the portal of the regional hydrological service of the Tuscany region (<https://www.sir.toscana.it/consistenza-rete>). The MT-CLIM mountain microclimate simulation model (Thornton & Running, 1999) was driven by temperature and precipitation series recorded at the nearby Vallombrosa meteorological station (Locality of Reggello, Florence; Lat 43.731° Lon 11.557; 980 m.a.s.l.) and was used to extrapolate this point measurement (referred to as the ‘base’ station) to the multiple inventory plot, making corrections for differences in elevation, slope and aspect between the base station and the given site. This dataset was utilized for historical simulations to validate the model.

To evaluate the performance of the 3D-CMCC-FEM model parameterization for *Pinus nigra* in the Vallombrosa forest, we based our analysis on two categories of variables: dendrometric variables (mean diameter at breast height (DBH), mean height (Ht), density (N)) and flux variables (Gross Primary Production (GPP) and Net Primary Production (NPP)). For flux variables, evaluation metrics were calculated on a monthly scale to capture seasonal variations. For dendrometric variables, evaluation metrics were assessed on annual scale to reflect their slower growth rates. This approach allows each variable to be analyzed at a time scale appropriate to its dynamics, ensuring also alignment with temporal resolution of available data. For each variable of interest, we performed a regression analysis between observed and simulated values to calculate the coefficient of determination (R^2^) and assess its significance. This metric provides insight into the strength of the linear relationship between observed and simulated data. We assessed ME (Nash-Sutcliffe model efficiency coefficient), normalized RMSE, and absolute bias AB_s_ for GPP, NPP, DBH and Ht on their respective temporal scale, where ME is calculated as: $1- \frac{\sum_{1}^{n} {(O_{i}-P_{i})}^{2}}{\sum_{1}^{n} {(O_{i}-\bar{O})}^{2}}$, nRMSE as $\frac{\sqrt{\frac{1}{n} \sum_{1}^{n} {(O_{i}-P_{i})}^{2}}}{\bar{O}} \times100$ and AB_s_ as $\frac{1}{n} \sum_{1}^{n} \left| O_{i}-P_{i} \right|$.

where $O_{i}$  and $P_{i}$ are the observed and predicted values respectively, $\bar{O_{i}}$ is the mean of the observed values, and n is the number of observations.

The forest growth model demonstrates robust predictive performance, accurately capturing both the central tendency and variability of observed data across dendrometric and carbon flux variables (Table 1). For dendrometric variables, observed and predicted variables align closely: DBH, showed an observed mean of 26.55 ± 6.54 cm, closely matched by a predicted mean of 27.44 ± 7.85 cm. Ht had an observed mean of 17.80 ± 5.67 m and a predicted mean of 18.57 ± 4.94 m. Similarly, for N, the observed mean was 1450 ± 599 ha^−1^, compared to a predicted mean of 1332 ± 584 ha^−1^. Beyond mean and variance values, model performance across the dendrometric variables demonstrates a strong linear relationship between observed and predicted values for DBH and N, as reflected by R^2^ values of 0.90 and 0.87, respectively, both significant at p < 0.001. Height shows a moderate linear fit, with an R^2^ of 0.67, suggesting a slightly less precise fit for this variable. ME values further highlight the model’s effectiveness, with values close to 1 for DBH (0.82) and N (0.83), indicating that the model’s predictions for these variables surpass the baseline of using the mean of the observed data. Height shows an ME of 0.65, indicating acceptable predictive performance and reasonable alignment with observed data. NRMSE values provide insight into prediction error, with low values for DBH (0.13) and N (0.11), indicating minimal deviation between observed and predicted values relative to the range of observed values. Height has a slightly higher NRMSE of 0.15, suggesting a modestly greater proportion of error for this variable. The AB_s_ reveals a slight overestimation for DBH (0.13) and Ht (0.14) and a minor underestimation for N (-0.19).

For carbon flux variables, the model similarly captures the observed data’s central tendency and variability. GPP showed an observed mean was 109.86 ± 60.61 gC.m^-2^.month^-1^, closely matched by a predicted mean of 114.62 ± 67.96 gC.m^-2^.month^-1^. Similarly, NPP exhibited an observed mean of 73.95 ± 54.83 gC.m^-2^.month^-1^ and a predicted mean of 66.19 ± 42.60 gC.m^-2^.month^-1^, indicating that the model captures the central tendency and variability of observed data for both variables. The R^2^ values further confirm the model’s predictive accuracy, with GPP showing a strong linear relationship between observed and predicted values (R^2^ = 0.87, p<0.001). NPP exhibits a moderate fit (R^2^= 0.64, p<0.001), reflecting reasonable predictive accuracy, albeit slightly lower than GPP. ME scores are high for GPP (0.86), indicating substantial improvement over the mean of observed data, while NPP shows an ME of 0.62, suggesting acceptable predictive performance. NRMSE values provide insight into prediction error, with GPP showing a low NRMSE of 0.11, indicating minimal deviation relative to the observed range, and NPP displaying a slightly higher NRMSE of 0.13, suggesting a modest increase in error proportion. The AB_s_ reveals a slight overestimation for GPP (0.07) and a minor underestimation for NPP (-0.14), indicating minimal systematic bias in model predictions.

Table 1: Observed and predicted means (± standard deviation) and evaluation metrics for model performance in predicting annual dendrometric variables (DBH (cm), Ht (m), N (ha^-1^)) and monthly flux variables (GPP and NPP; in gC.m^-2^.month^-1^). *R² values are significant at *p < 0.001*.

|  | **Dendrometric variable** | | | **Flux variable** | |
| --- | --- | --- | --- | --- | --- |
| **Metric** | **DBH** | **Ht** | **N** | **GPP** | **NPP** |
| **Observed ± SD** | 26.55 ± 6.54 | 17.80 ± 5.67 | 1450 ± 599 | 109.86 ± 60.61 | 73.95 ± 54.83 |
| **Predicted ± SD** | 27.44 ± 7.85 | 18.57 ± 4.94 | 1332 ± 584 | 114.62 ± 67.96 | 66.19 ± 42.60 |
| **R^2^** | 0.90* | 0.67* | 0.87* | 0.87* | 0.64* |
| **ME** | 0.82 | 0.65 | 0.83 | 0.86 | 0.62 |
| **NRMSE** | 0.12 | 0.15 | 0.11 | 0.11 | 0.13 |
| **AB_S_** | 0.13 | 0.14 | -0.19 | 0.07 | -0.14 |

Table 2: Bucking allocation criteria according to different sectors (Cutting started from 30 cm above ground level)

| **Product** | **Priority** | **Number** | **Length** | **Diameter** | **Quality criteria** |
| --- | --- | --- | --- | --- | --- |
| Stump | 1 | 1 | 0.3 m |  | - |
| Interior carpentry | 2 | - | 3 – 5 m | Ø_SE_ ≥ 25 cm | (Ø_KW_ / Ø_HW_)^2^ ≤ 13% |
| Lumber | 3 | - | 3 – 6 m | Ø_SE_ ≥ 25 cm | Ø_KW_/ Ø_SE_ ≤ 30 % |
| Sawing | 4 | - | 3 – 5 m | Ø_SE_ ≥ 25 cm | - |
| Particle boards | 5 | 3 | 2 – 2.2 m | Ø_SE_ ≥ 7 cm | - |
| Paper pulp | 6 | - | 2.4 – 4.8 | Ø_SE_ ≥ 7 cm | - |
| Firewood | 7 | - | 0.5 – 1 m | Ø_SE_ ≥ 7 cm | - |
| Toplog | 8 | - | - |  | - |

Ø_SE_: Diameter small end, Ø_KW_: Diameter of knotted wood, Ø_HW_: Diameter of Heartwood

Table 3: Efficiency and loss of different wood products reported to the log volume including bark.

| **Product** | **Efficiency (%)** | **Loss (%)** |
| --- | --- | --- |
| Stump | 1 | 0 |
| Furniture | 0.45 | 0.55 |
| Lumber | 0.50 | 0.50 |
| Sawing | 0.40 | 0.60 |
| Particle boards | 0.76 | 0.24 |
| Paper pulp | 0.8 | 0.20 |
| Firewood | 1 | 0 |
| TopLog | 1 | 0 |

Table 4: Allocation of the harvested wood product losses.

|  | **Fire (%)** | **Boards (%)** | **Paper (%)** | **Millsite (%)** |
| --- | --- | --- | --- | --- |
| **Furniture** | 0.15 | 0.1 | 0.2 | 0.1 |
| **Lumber** | 0.15 | 0.1 | 0.15 | 0.1 |
| **Sawing** | 0.2 | 0.2 | 0 | 0.1 |
| **Particle boards** | 0.14 | 0 | 0 | 0.2 |
| **Paper pulp** | 0 | 0 | 0 | 0 |

Table 5: Lifespan and recycling rate of different wood products and disposal.

| **Wood product** | **Average lifespan (yrs.)** | **Recycling rate (%)** |
| --- | --- | --- |
| Furniture | 30 | 0.1 |
| Lumber | 65 | 0.1 |
| Sawn wood | 50 | 0.1 |
| Particle boards | 20 | 0.1 |
| Paper pulp | 1 | 0.1 |
| Firewood including the BGB, stump, and the Top Log | 1 | - |
| Mill site dump | 5 | - |
| Landfill | 145 | - |

Table 6: Assumed substitution between wood products and substitutes in the considered services. The fraction of product in the likely substitution scenario and the average reduction in CC impacts (aka. displacement factor) were documented from (Sathre & O’Connor, 2010; Suter et al., 2017).

| **Wood product** | **Substitute** | **Service** | **Likely fraction of product in the substitution scenario (%)** | **The average reduction in CC impacts (tCO2-eq.m^-3^)** | **Displacement factor (tCO2-eq.m^-3^)** |
| --- | --- | --- | --- | --- | --- |
| Furniture | Polypropylene | Furniture | 47 | 0.61 | 0.82 |
|  | Steel, chromium, secondary | Furniture | 47 | 0.99 |  |
|  | Steel, chromium, primary | Furniture | 6 | 1.23 |  |
| Lumber | Steel, secondary | Construction | 70 | -0.06 | 0.06 |
|  | Steel, primary | Construction | 10 | 0.58 |  |
|  | Concrete | Construction | 10 | 0.27 |  |
|  | Brick | Construction | 10 | 0.18 |  |
| Sawn wood | Concrete | Construction | 39 | 0.27 | 0.32 |
|  | Brick | Construction | 31 | 0.18 |  |
|  | Polyethylene | Packaging | 9 | 0.30 |  |
|  | Aluminum, secondary | Packaging | 0.9 | 0.09 |  |
|  | Aluminum, primary | Packaging | 0.1 | 3.67 |  |
|  | Polypropylene | Furniture | 10 | 0.30 |  |
|  | Steel, chromium, secondary | Furniture | 9 | 0.99 |  |
|  | Steel, chromium, primary | Furniture | 1 | 1.23 |  |
| Particle boards | Plaster | Construction | 60 | 0.06 | 0.2 |
|  | Glass | Furniture | 40 | 0.41 |  |
| Paper pulp | None | Print | 100 | - | - |
| Firewood including the TopLog | Natural gas | Energy | 30 | 0.32 | 0.48 |
|  | Fuel oil | Energy | 70 | 0.55 |  |

PS: The volume in the displacement factor refers to the dry volume.

Table 7: 3D-CMCC-FEM model paramterization

| **Parameter** | **Value** | **Species** | **Reference** | **Description** | **Citation** |
| --- | --- | --- | --- | --- | --- |
| LIGHT_TOL | 4 | P. nigra | Isajev et al., 2003 | Light tolerance: 4 = very shade intolerant (cc = 90%), 3 = shade intolerant (cc = 100%), 2 = shade tolerant (cc = 110%), 1 = very shade tolerant (cc = 120%) (dimensionless) (cc = canopy cover) | Isajev, V.; Fady, B.; Semerci, H.; Andonovski, V. (2003) European black pine (Pinus nigra). EUFORGEN Technical Guidelines for Genetic Conservation and Use 6 p. ISBN: 978-92-9043-659-1, ISBN: 92-9043-659-X |
| PHENOLOGY | 1.2 | P. nigra | Evidence knowledge | Phenology of species: 0.1 = deciduous broadleaf, 0.2 = deciduous needleleaf, 1.1 = broadleaf evergreen, 1.2 = needleleaf evergreen (dimensionless) |  |
| K | 0.5 | P. nigra | Navarro-Cerrillo et al., 2016 | Extinction coefficient for absorption of PAR by canopy (ratio) | [http://dx.doi.org/10.5424/fs/2016253-08610.](http://dx.doi.org/10.5424/fs/2016253-08610) |
| ALBEDO | 0.116 | P. sylvestris | Calculated from Kuusinen et al., 2014 | Canopy albedo (ratio) | https://doi.org/10.1016/j.ecolmodel.2014.04.007. |
| SLA_AVG0 | 5 | P. nigra | Navarro-Cerrillo et al., 2016 | Average specific leaf area (juvenile) (m2 / Kg DM) | [http://dx.doi.org/10.5424/fs/2016253-08610.](http://dx.doi.org/10.5424/fs/2016253-08610) |
| SLA_AVG1 | 4 | P. nigra | Navarro-Cerrillo et al., 2016 | Average specific leaf area (mature) (m2 / Kg DM) | [http://dx.doi.org/10.5424/fs/2016253-08610.](http://dx.doi.org/10.5424/fs/2016253-08610) |
| TSLA | 4 | P. nigra | Navarro-Cerrillo et al., 2016 | Age at which SLA_AVG = (SLA_AVG0 + SLA_AVG1) / 2 (yr) | http://dx.doi.org/10.5424/fs/2016253-08610. |
| SLA_RATIO | 2 | P. pinaster Aiton | Nunes et al., 2014 | Ratio of shaded to sunlit projected SLA (ratio) | https://doi.org/10.1093/forestry/cpu044 |
| LAI_RATIO | 2.6 | P. pinaster Aiton | Nunes et al., 2014 | Ratio of all-sided to projected leaf area (ratio) | https://doi.org/10.1093/forestry/cpu044 |
| FRACBB0 | 0.5 | P. nigra | Navarro-Cerrillo et al., 2016 | Branch and bark fraction (juvenile) (ratio) | http://dx.doi.org/10.5424/fs/2016253-08610. |
| FRACBB1 | 0.1 | P. nigra | Navarro-Cerrillo et al., 2016 | Branch and bark fraction (mature) (ratio) | http://dx.doi.org/10.5424/fs/2016253-08610. |
| TBB | 5 | P. nigra | Navarro-Cerrillo et al., 2016 | Age at which FRACBB = (FRACBB0 + FRACBB1 ) / 2 (yr) | http://dx.doi.org/10.5424/fs/2016253-08610. |
| RHO0 | 0.43 | P. nigra | Navarro-Cerrillo et al., 2016 | Minimum basic density (juvenile) (Mg DM / m3) | http://dx.doi.org/10.5424/fs/2016253-08610. |
| RHO1 | 0.43 | P. nigra | Navarro-Cerrillo et al., 2016 | Maximum basic density (mature) (Mg DM / m3) | http://dx.doi.org/10.5424/fs/2016253-08610. |
| TRHO | 4 | P. nigra | Navarro-Cerrillo et al., 2016 | Age at which RHO = (RHO0 + RHO1) / 2 (yr) | http://dx.doi.org/10.5424/fs/2016253-08610. |
| COEFFCOND | 0.05 | P. nigra | Navarro-Cerrillo et al., 2016 | Controls stomatal response to VPD (mbar) | http://dx.doi.org/10.5424/fs/2016253-08610. |
| BLCOND | 0.2 | P. nigra | Navarro-Cerrillo et al., 2016 | Canopy boundary layer conductance (m / s) | http://dx.doi.org/10.5424/fs/2016253-08610. |
| MAXCOND | 0.018 | P. nigra | Navarro-Cerrillo et al., 2016 | Maximum leaf (stomatal) conductance (m / s) | http://dx.doi.org/10.5424/fs/2016253-08610. |
| CUTCOND | 0.00006 | P. sylvestris | Tatarinov et al., 2006 | Cuticular conductance (m / s) | https://doi.org/10.1016/j.foreco.2006.09.085 |
| MAXAGE | 461 | P. nigra | Nagel & Cerioni 2023 | Controls rate of physiological decline of forest (yr) | https://doi.org/10.1007/s10342-023-01540-5 |
| RAGE | 0.95 | P. nigra | Navarro-Cerrillo et al., 2016 | Relative age to give f(AGE) = 0.5 (dimensionless) | https://doi.org/10.1093/forestry/cpu044 |
| NAGE | 4 | P. nigra | Navarro-Cerrillo et al., 2016 | Power of relative age in f(AGE) (dimensionless) | http://dx.doi.org/10.5424/fs/2016253-08610. |
| GROWTHTMIN | 0 | P. nigra | Navarro-Cerrillo et al., 2016 | Minimum temperature for growth (°C) | http://dx.doi.org/10.5424/fs/2016253-08610. |
| GROWTHTMAX | 35 | P. nigra | Navarro-Cerrillo et al., 2016 | Maximum temperature for growth (°C) | http://dx.doi.org/10.5424/fs/2016253-08610. |
| GROWTHTOPT | 15 | P. nigra | Navarro-Cerrillo et al., 2016 | Optimum temperature for growth (°C) | http://dx.doi.org/10.5424/fs/2016253-08610. |
| SWPOPEN | -0.65 | P. pinaster Aiton | Nunes et al., 2014 | Leaf water potential (start of reduction) (MPa) | https://doi.org/10.1093/forestry/cpu044 |
| SWPCLOSE | -2.5 | P. pinaster Aiton | Nunes et al., 2014 | Leaf water potential (end of reduction) (Mpa) | https://doi.org/10.1093/forestry/cpu044 |
| OMEGA | 0.5 | Needle-leaf | Arora and Boer, 2004 | Controls sensibility of allocation to changes in water and light availability (dimensionless) | https://doi.org/10.1111/j.1365-2486.2004.00890.x |
| S0 | 0.74 | P. sylvestris | Chmura et al., 2012 | Stem allocation factor (ratio) | http://dx.doi.org/10.1080/02827581.2013.844269 |
| R0 | 0.16 | P. sylvestris | Chmura et al., 2012 | Root allocation factor (ratio) | http://dx.doi.org/10.1080/02827581.2013.844269 |
| F0 | 0.1 | P. sylvestris | Chmura et al., 2012 | Foliage allocation factor (ratio) | http://dx.doi.org/10.1080/02827581.2013.844269 |
| FRUIT_PERC | 6.17 | P. nigra | Fernández-Martínez et al., 2016 | Fraction of NPP allocated for reproduction during the prescribed seasonal period (ratio) | https://doi.org/10.1111/nph.14193 |
| FINE_ROOT_LEAF | 1 | P. sylvestris | Tatarinov et al., 2006 | Fine root C:leaf C (ratio) | https://doi.org/10.1016/j.foreco.2006.09.085 |
| COARSE_ROOT_STEM | 0.44 | P. sylvestris | Tatarinov et al., 2006 | Coarse root C:stem C (ratio) | https://doi.org/10.1016/j.foreco.2006.09.085 |
| LIVE_TOTAL_WOOD | 0.076 | P. sylvestris | Tatarinov et al., 2006 | Live C:total wood C (ratio) | https://doi.org/10.1016/j.foreco.2006.09.085 |
| N_RUBISCO | 0.055 | P. sylvestris | Tatarinov et al., 2006 | Fraction of leaf N in Rubisco (ratio) | https://doi.org/10.1016/j.foreco.2006.09.085 |
| CN_LEAVES | 36 | P. sylvestris | Tatarinov et al., 2006 | C:N of leaves (ratio) | https://doi.org/10.1016/j.foreco.2006.09.085 |
| CN_FINE_ROOTS | 49 | P. sylvestris | Tatarinov et al., 2006 | C:N of fine roots (ratio) | https://doi.org/10.1016/j.foreco.2006.09.085 |
| CN_LIVEWOOD | 58 | P. sylvestris | Tatarinov et al., 2006 | C:N of live woods (ratio) | https://doi.org/10.1016/j.foreco.2006.09.085 |
| LEAF_FINERROOT_TURNOVER | leaf: 0.39 / fine roots : 0.868 | P. sylvestris | Tatarinov et al., 2006 | Average annual leaf and fine root turnover (yr–1) | https://doi.org/10.1016/j.foreco.2006.09.085 |
| LIVEWOOD_TURNOVER | 0.02 |  | FIXED | Annual livewood turnover (yr^–1^) |  |
| SAPWOOD_TURNOVER | 0.02 |  | FIXED | Annual sapwood turnover (yr^–1^) |  |
| DBHDCMIN | 0.12 | P. nigra | Estimated from plot data | Minimum DBH to crown diameter (ratio) |  |
| SAP_A | 0.36 | P. ponderosa | Mitra et al., 2019 | Scaling coefficient in sapwood area to DBH relationship (dimensionless) | https://doi.org/10.1007/s11676-019-01048-y |
| SAP_B | 2.08 | P. ponderosa | Mitra et al., 2019 | Scaling coefficient in sapwood area to DBH relationship (exp) (dimensionless) | https://doi.org/10.1007/s11676-019-01048-y |
| SAP_LEAF | 1525 | P. sylvestris | Calculated from : Mencuccini and Grace (1996), Van Hees and Bartelink (1993), Vanninen et al. (1996), and Albrekston (1984) | Leaf area to sapwood area (ratio) | [https://doi.org/10.1093/treephys/15.1.1 / https://doi.org/10.1016/0378-1127(93)90128-A / https://doi.org/10.1007/BF02185674 / https://doi.org/10.1093/forestry/57.1.35](https://doi.org/10.1093/treephys/15.1.1%20-) |
| SAP_WRES | 0.05 | Evergreen | Evidence knowledge | Sapwood to reserve biomass: 0.11 = deciduous, 0.05 = evergreen (ratio) |  |
| STEMCONST_P | 0.20374 | P. nigra | Estimated from plot data | Scaling coefficient in stem mass to DBH relationship (dimensionless) |  |
| STEMPOWER_P | 2.197 | P. nigra | Estimated from plot data | Scaling coefficient in stem mass to DBH relationship (exp) (dimensionless) |  |
| CRA | 33.67 | P. nigra | Estimated from plot data | Chapman-Richards asymptotic maximum height (m) |  |
| CRB | 0.1 | P. nigra | Estimated from plot data | Chapman-Richards exponential decay parameter (dimensionless) |  |
| CRC | 6.5 | P. nigra | Estimated from plot data | Chapman-Richards shape parameter (dimensionless) |  |
| CROWN_A | 1.277 | P. nigra | Estimated from plot data | Scaling coefficient in crown length to height relationship (dimensionless) |  |
| CROWN_B | 0.589 | P. nigra | Estimated from plot data | Scaling coefficient in crown length to height relationship (exp) (dimensionless) |  |
| SEXAGE | 25 | P. nigra | Calculated from Debain et al., 2007 | Age at sexual maturity (yr) | https://doi.org/10.1139/X06-265 |
