## Supplementary Material 2 for "Long-term forest-sector mitigation and radiative forcing under contrasting management, climate, and substitution pathways"

**Supplementary 2: Supporting Analyses**

**Statistical Modeling details**

**Linear Mixed-Effects Model (LME)**

LME were used to analyze gross primary productivity (GPP), ecosystem respiration (RECO), net ecosystem exchange (NEE), and forest sector balance (FSB). Models were implemented using the lme4 R package. Fixed effects varied by response variable and included combinations of Climate, Management, Century, Wood Use (WU), and Substitution (SUB). YEAR was included as a random intercept to account for temporal autocorrelation.

Residual diagnostics revealed violations of model assumptions in several cases. To address these, response variables were transformed using the Yeo-Johnson method via the bestNormalize package. Influential outliers were identified and removed based on Cook’s Distance, using the car package. Two-way interactions among predictors were tested, and statistical significance was assessed using Type III ANOVA with Satterthwaite’s approximation (lmerTest package). Effect sizes were interpreted using partial eta-squared values, calculated with the effectsize package, following Cohen’s (1988) thresholds for small (η² ≈ 0.01), medium (η² ≈ 0.06), and large (η² ≥ 0.14) effects.

Response ∼ Climate + Management + Century +

                    Climate × Management +

                    Climate × Century +

                    Management × Century

FSB ∼ Climate + Management + WU + SUB +

            Climate × Century +

            Management × Century +

            Climate × SUB +

            Management × SUB +

            Management × Climate

**Generalized Additive Mixed Models (GAMMs)**

Generalized additive mixed models (GAMMs) were applied to model wood emissions, substitution effects (material, energy, and their sum), and wood balance (Wbalance). These models were implemented using the mgcv package. Fixed effects included Climate, Management, Wood Use (WU), and Substitution (SUB). A smooth term was added for Century to capture long-term non-linear trends:

s(Century, k = 3)

Two-way interactions (e.g., Climate × SUB, Management × SUB, Management × Climate) were included to assess joint effects. Final model selection was based on Akaike Information Criterion (AIC) and likelihood ratio tests (LRTs).

The final model structure was:

Response ∼ Climate + Management + WU + SUB +

                   Climate × SUB +

                   Management × SUB +

                   Management × Climate +

                   s(Century)

**Table 1:** Century-scale dynamics of ecosystem fluxes by forest management and climate scenario (unit: gC m^-2^ yr^-1^)

|  | | **21st century** | | | **22nd century** | | | **23rd century** | | | **All** | | |
| --- | --- | --- | --- | --- | --- | --- | --- | --- | --- | --- | --- | --- | --- |
| **Management** | **Climate** | **GPP** | **RECO** | **NEE** | **GPP** | **RECO** | **NEE** | **GPP** | **RECO** | **NEE** | **GPP** | **RECO** | **NEE** |
| TM | CUR | 405 | 299 | -106 | 537 | 365 | -172 | 510 | 353 | -156 | 510 | 352 | -157 |
| TM | SSP1-2.6 | 476 | 372 | -103 | 629 | 428 | -200 | 613 | 421 | -191 | 605 | 419 | -185 |
| TM | SSP5-8.5 | 557 | 432 | -124 | 597 | 465 | -131 | 232 | 228 | -3 | 507 | 405 | -101 |
| ADAPT | CUR | 266 | 217 | -49 | 249 | 186 | -62 | 296 | 205 | -91 | 271 | 202 | -68 |
| ADAPT | SSP1-2.6 | 332 | 280 | -52 | 325 | 241 | -84 | 365 | 255 | -109 | 341 | 258 | -83 |
| ADAPT | SSP5-8.5 | 411 | 336 | -75 | 190 | 182 | -8 | 46 | 49 | 2 | 205 | 180 | -24 |
| BIOE | CUR | 362 | 273 | -89 | 487 | 345 | -142 | 588 | 404 | -183 | 485 | 344 | -140 |
| BIOE | SSP1-2.6 | 442 | 351 | -91 | 557 | 401 | -156 | 653 | 461 | -191 | 557 | 407 | -149 |
| BIOE | SSP5-8.5 | 537 | 420 | -116 | 476 | 385 | -91 | 111 | 101 | -10 | 366 | 295 | -70 |
| TRANS | CUR | 450 | 348 | -102 | 706 | 626 | -79 | 673 | 635 | -38 | 616 | 545 | -70 |
| TRANS | SSP1-2.6 | 518 | 427 | -90 | 867 | 744 | -123 | 838 | 770 | -68 | 796 | 706 | -90 |
| TRANS | SSP5-8.5 | 613 | 504 | -109 | 658 | 712 | 53 | 164 | 247 | 83 | 476 | 487 | 10 |
| WOOD | CUR | 296 | 233 | -63 | 358 | 247 | -110 | 333 | 253 | -79 | 331 | 245 | -85 |
| WOOD | SSP1-2.6 | 372 | 302 | -70 | 481 | 333 | -148 | 426 | 328 | -98 | 429 | 322 | -107 |
| WOOD | SSP5-8.5 | 464 | 365 | -99 | 352 | 293 | -58 | 72 | 73 | 1 | 286 | 237 | -49 |

Table 2: Summary of linear mixed-effects results for GPP, RECO, and NEE. The table reports the degree of freedom (Df), Chi-square test statistic (Chisq), p-value (Pr>Chisq), and effect size (η²) for fixed effects and their interactions.

| **Response** | **Variable** | **Df** | **Chisq** | **Pr(>Chisq)** | **η²** |
| --- | --- | --- | --- | --- | --- |
| **GPP** | (Intercept) | 1 | 496.866 | <0.001 |  |
|  | Climate | 2 | 285.547 | <0.001 | 0.288 |
|  | Management | 4 | 202.039 | <0.001 | 0.219 |
|  | century | 2 | 13.150 | 0.001 | 0.071 |
|  | Climate:Management | 8 | 159.623 | <0.001 | 0.039 |
|  | Climate:century | 4 | 2,238.544 | <0.001 | 0.285 |
|  | Management:century | 8 | 359.242 | <0.001 | 0.061 |
| **NEE** | (Intercept) | 1 | 139.739 | <0.001 |  |
|  | Climate | 2 | 25.181 | <0.001 | 0.125 |
|  | Management | 4 | 97.078 | <0.001 | 0.115 |
|  | century | 2 | 6.964 | 0.031 | 0.030 |
|  | Climate:Management | 8 | 73.832 | <0.001 | 0.014 |
|  | Climate:century | 4 | 603.581 | <0.001 | 0.070 |
|  | Management:century | 8 | 318.960 | <0.001 | 0.026 |
| **RECO** | (Intercept) | 1 | 757.664 | <0.001 |  |
|  | Climate | 2 | 575.953 | <0.001 | 0.335 |
|  | Management | 4 | 321.655 | <0.001 | 0.520 |
|  | century | 2 | 14.242 | <0.001 | 0.107 |
|  | Climate:Management | 8 | 152.831 | <0.001 | 0.068 |
|  | Climate:century | 4 | 3,424.483 | <0.001 | 0.399 |
|  | Management:century | 8 | 1,286.206 | <0.001 | 0.162 |

Table 3: Model comparison results based on likelihood ratio tests and changes in AIC (ΔAIC) for fixed effects and interactions across six response variables: yearly emissions, yearly energy substitution, yearly material substitution, yearly total substitution, and wood balance. ΔAIC indicates the change in AIC when a term is removed from the full model, and the corresponding likelihood ratio test (LRT) p-value assesses statistical significance.

| **Response** | **Term** | **ΔAIC** | **LRT p-value** |
| --- | --- | --- | --- |
| **Emissions** | Management:SUB | -31.999 | 1.00000 |
|  | Climate:SUB | -16.000 | 1.00000 |
|  | WU | -5.609 | 0.94210 |
|  | Management | 0.000 | 0.00000 |
|  | Climate | 0.000 | 0.00000 |
|  | SUB | 0.000 | 1.00000 |
|  | Management:Climate | 83.824 | 0.00000 |
| **Energy substitution** | WU | -5.853 | 0.98570 |
|  | Management:Climate | -5.315 | 0.22020 |
|  | Management | 0.000 | 1.00000 |
|  | Climate | 0.000 | 0.00000 |
|  | SUB | 0.000 | 0.00000 |
|  | Climate:SUB | 3.150 | 0.01408 |
|  | Management:SUB | 98.361 | 0.00000 |
| **Material substitution** | WU | -5.948 | 0.99690 |
|  | Management:Climate | -3.391 | 0.12600 |
|  | Management | 0.000 | 0.00000 |
|  | Climate | 0.000 | 0.00000 |
|  | SUB | 0.000 | 0.00000 |
|  | Climate:SUB | 4.377 | 0.00900 |
|  | Management:SUB | 87.145 | 0.00000 |
| **Total substitution** | WU | -5.925 | 0.99470 |
|  | Management:Climate | -3.037 | 0.11310 |
|  | Management | 0.000 | 0.00000 |
|  | Climate | 0.000 | 0.00000 |
|  | SUB | 0.000 | 0.00000 |
|  | Climate:SUB | 6.454 | 0.00414 |
|  | Management:SUB | 101.446 | 0.00000 |
| **Wood balance** | Climate:SUB | -10.207 | 0.67040 |
|  | WU | -5.751 | 0.96940 |
|  | Management | 0.000 | 1.00000 |
|  | Climate | 0.000 | 1.00000 |
|  | SUB | 0.000 | 0.00000 |
|  | Management:SUB | 3.952 | 0.00294 |
|  | Management:Climate | 57.097 | 0.00000 |

Table 4: Summary of LME results: Chi-squared, p-values, and partial η² for each response variable.

| **Response** | **Variable** | **df** | **F-value** | **Pr(>F-value)** | **η²** |
| --- | --- | --- | --- | --- | --- |
| **FSB** | Climate | 2 | 21827.699 | <0.001 | 0.43 |
|  | Management | 4 | 3067.448 | <0.001 | 0.18 |
|  | Century | 2 | 6.215 | 0.002 | 0.04 |
|  | Substitution | 4 | 485.861 | <0.001 | 0.03 |
|  | Climate:Century | 4 | 8366.336 | <0.001 | 0.37 |
|  | Management:Century | 8 | 1347.196 | <0.001 | 0.16 |
|  | Management:Climate | 8 | 573.072 | <0.001 | 0.07 |
|  | Climate:Substitution | 8 | 10.226 | <0.001 | 0.0001 |
|  | Management:Substitution | 16 | 22.174 | <0.001 | 0.0006 |

Table 5: Contribution percentage of every component to FSB by decarbonization scenario and management scheme

| **Variable** | **Factor** | **NEE** | **Emissions** | **Substitution** |
| --- | --- | --- | --- | --- |
| Substitution | SUB0 | 0.53 ± 0.04 | 0.29 ± 0.02 | 0.18 ± 0.03 |
|  | SUB25 | 0.60 ± 0.04 | 0.33 ± 0.02 | 0.07 ± 0.01 |
|  | SUB50 | 0.64 ± 0.03 | 0.34 ± 0.03 | 0.02 ± 0.00 |
|  | SUB75 | 0.65 ± 0.03 | 0.35 ± 0.03 | 0.00 ± 0.00 |
|  | SUB100 | 0.65 ± 0.03 | 0.35 ± 0.03 | 0.00 ± 0.00 |
| Management | ADAPT | 0.60 ± 0.05 | 0.34 ± 0.03 | 0.06 ± 0.08 |
|  | BIOE | 0.61 ± 0.05 | 0.33 ± 0.03 | 0.06 ± 0.08 |
|  | TM | 0.67 ± 0.04 | 0.29 ± 0.02 | 0.04 ± 0.05 |
|  | TRANS | 0.57 ± 0.05 | 0.37 ± 0.03 | 0.06 ± 0.05 |
|  | WOOD | 0.60 ± 0.05 | 0.34 ± 0.03 | 0.06 ± 0.05 |

Table 6: Evolution of harvested wood volume (m^3^ ha^-1^ yr^-1^) across management strategies, climate scenarios, and centuries

| **Management** | **Climate** | **21st century** | **22nd century** | **23rd century** | **Full period** | **Comments** |
| --- | --- | --- | --- | --- | --- | --- |
| ADAPT | CUR | 1.66 | 3.38 | 3.05 | 2.76 | ADAPT management scenario is a 100-year rotation managed following a clearcut silviculture. This implies that harvest occurs two times during the study period (2115 and 2215). Under the CUR and SSP1-2.6 scenarios, harvesting occurs mainly in the second and third centuries retrieving equal volumes. Under the SSP5-8.5, however, harvested volume was almost null by the last century. |
|  | SSP1-2.6 | 2.06 | 4.72 | 3.96 | 3.65 |  |
|  | SSP5-8.5 | 2.70 | 4.37 | 0.29 | 2.43 |  |
| BIOE | CUR | 6.21 | 5.70 | 6.69 | 6.20 | BIOE management scenario is a 80-year rotation managed following shelterwood principles. This implies that harvest (final cut) occurs three times during the study period (2095, 2175, and 2255). Under the CUR and SSP1-2.6 scenarios, harvesting occurs in each century, retrieving almost equal volumes. Under the SSP5-8.5, however, harvested volume was almost null by the last century. |
|  | SSP1-.26 | 7.68 | 6.53 | 7.25 | 7.10 |  |
|  | SSP5-8.5 | 9.54 | 4.47 | 0.28 | 4.48 |  |
| TM | CUR | 1.86 | 5.94 | 5.29 | 4.51 | TM management scenario is a selective system where cutting occurs every 5-years with the aim of harvesting 20% of the standing volume under the condition of maintaining a minimum growing stock (GSV) of 120 m^3^. Across all climate scenarios during the first century, thinning is moderate as the algorithm waits for the condition of GSV = 120 m3 to be met. Under CUR and SSP1-2.6, harvested volume is of the same magnitude during the second and third century. Under the SSP5-8.5 scenario, in the third century, volume tends to be null. |
|  | SSP1-2.6 | 2.43 | 7.25 | 6.27 | 5.46 |  |
|  | SSP5-8.5 | 3.36 | 6.61 | 0.62 | 3.53 |  |
| TRANS | CUR | 1.57 | 3.45 | 0.00 | 1.68 | TRANS management is a scenario that aims to gradually transition from a forest managed under an even-aged structure to an unmanaged forest. 2150 represents the year of last intervention, which means that no harvesting occurs thereafter. Most of the volume was harvested during the second century. |
|  | SSP1-2.6 | 1.77 | 5.06 | 0.00 | 2.30 |  |
|  | SSP5-8.5 | 2.31 | 7.40 | 0.00 | 3.28 |  |
| WOOD | CUR | 0.86 | 5.55 | 3.89 | 3.48 | WOOD management scenario is a 130-year rotation managed following shelterwood principles. This implies that harvest (final cut) occurs two times during the study period (2145 and 2275). During the second century, harvested volume reaches its maximum under the different climate scenarios, then it declines during the last century under CUR and SSP1-2.6 while it reaches nullity under SSP5-8.5. |
|  | SSP1-2.6 | 0.98 | 8.30 | 5.58 | 5.00 |  |
|  | SSP5-8.5 | 1.31 | 10.53 | 0.27 | 3.85 |  |

Table 7: Summary of harvested tree counts, volume, and DBH distribution by management and climate scenario. DBH metrics (in cm) include minimum, quartiles (Q1, Q3), median, mean, maximum, and standard deviation (SD). "PctAbove30" indicates the percentage of trees with DBH above 30 cm. “Nabove30” indicates the number of trees with DBH above 30 cm.

| **Management** | **Climate** | **Harvested trees (ha^-1^)** | **Harvested volume (m^3^)** | **Min (cm)** | **Q1 (cm)** | **Median (cm)** | **Mean (cm)** | **Q3 (cm)** | **Max (cm)** | **SD (cm)** | **PctAbove30 (%)** | **Nabove30** |
| --- | --- | --- | --- | --- | --- | --- | --- | --- | --- | --- | --- | --- |
| TM | CUR | 8960 | 1279 | 14.12 | 19.86 | 21.29 | 21.81 | 23.29 | 44.11 | 4.41 | 4.80 | 430 |
| TM | SSP1-2.6 | 10070 | 1556 | 14.20 | 19.69 | 21.36 | 22.08 | 23.21 | 56.23 | 5.30 | 4.55 | 458 |
| TM | SSP5-8.5 | 5078 | 1005 | 12.83 | 18.46 | 19.62 | 22.42 | 23.72 | 73.09 | 8.73 | 11.22 | 558 |
| ADAPT | CUR | 8934 | 782 | 3.02 | 3.03 | 3.03 | 10.37 | 19.55 | 43.68 | 11.55 | 11.08 | 1054 |
| ADAPT | SSP1-2.6 | 8904 | 1041 | 2.93 | 3.07 | 3.13 | 11.14 | 20.54 | 50.70 | 13.03 | 13.59 | 1210 |
| ADAPT | SSP5-8.5 | 9020 | 692 | 2.15 | 2.96 | 3.02 | 8.43 | 9.97 | 56.34 | 11.10 | 5.61 | 505 |
| BIOE | CUR | 19846 | 1761 | 3.13 | 4.79 | 5.03 | 11.07 | 10.98 | 45.37 | 11.00 | 14.00 | 2778 |
| BIOE | SSP1-2.6 | 19850 | 2022 | 3.04 | 4.72 | 5.10 | 11.38 | 11.09 | 49.96 | 11.72 | 14.05 | 2779 |
| BIOE | SSP5-8.5 | 20040 | 1276 | 3.08 | 3.11 | 5.05 | 8.47 | 8.77 | 54.37 | 10.04 | 5.45 | 1092 |
| TRANS | CUR | 3142 | 476 | 7.72 | 7.72 | 16.88 | 19.04 | 25.52 | 48.07 | 9.81 | 11.33 | 355 |
| TRANS | SSP1-2.6 | 3132 | 654 | 7.54 | 7.54 | 17.07 | 19.95 | 22.58 | 56.30 | 12.37 | 12.20 | 375 |
| TRANS | SSP5-8.5 | 2518 | 934 | 7.77 | 7.77 | 18.85 | 22.34 | 32.06 | 85.84 | 17.78 | 28.67 | 705 |
| WOOD | CUR | 12376 | 938 | 6.41 | 6.41 | 6.51 | 11.44 | 12.14 | 61.47 | 9.62 | 6.33 | 783 |
| WOOD | SSP1-2.6 | 12340 | 1354 | 6.50 | 6.50 | 6.63 | 12.24 | 12.42 | 74.97 | 11.57 | 6.43 | 793 |
| WOOD | SSP5-8.5 | 12432 | 1042 | 3.73 | 3.73 | 4.58 | 8.64 | 7.76 | 83.81 | 11.12 | 3.39 | 421 |

Table 8: Generalized linear model (GLM) coefficients for material and energy substitution responses, showing the effects of maximum DBH (maxDBH), the number of trees above 30 cm DBH (NDBH30), and volume (see colab mat|sub)*.*

| **Response** | **Term** | **Coefficient** | **std** | **z** | **P>\|z\|** |
| --- | --- | --- | --- | --- | --- |
| Material substitution  (R^2^= 0.96) | Intercept | -0.0197 | 0.012 | -1.606 | 0.108 |
|  | maxDBH | 0.0006 | 0.000 | 3.815 | 0.000 |
|  | NDBH30 | 1.644e-05 | 3.85e-06 | 4.273 | 0.000 |
|  | volume | 0.0019 | 0.002 | 1.170 | 0.242 |
| Energy substitution  (R^2^= 0.81) | Intercept | 0.0013 | 0.012 | 0.107 | 0.915 |
|  | maxDBH | 0.0003 | 0.000 | 1.793 | 0.073 |
|  | NDBH30 | 3.521e-06 | 3.87e-06 | 0.910 | 0.363 |
|  | volume | 0.0051 | 0.002 | 3.099 | 0.002 |

Table 9: Summary of linear regression results for RFbio vs.CB

| **Term** | **Estimate** | **Std_error** | **t-value** | **Pr(>\|t\|)** |
| --- | --- | --- | --- | --- |
| Intercept | 4.979 | 0.921 | 5.403 | 1.75e-07 |
| total_CM | 0.441 | 0.006 | 65.817 | <2e-16 |

*Model fit statistics*: (*R^2^ = 0.9531; p-value < 2.2×10^-16^)*

Table 10: Mitigation efficiency of forest sector under various management schemes (ADAPT, BIOE, TM, TRANS, and WOOD) climate scenarios (CUR, SSP1-2.6, and SSP5-8.5), decarbonization pathways (SUB0-SUB100) and time horizons (H1, H2, and H3).

|  | | **ADAPT** | | | **BIOE** | | | **TM** | | | **TRANS** | | | **WOOD** | | |
| --- | --- | --- | --- | --- | --- | --- | --- | --- | --- | --- | --- | --- | --- | --- | --- | --- |
| **Climate** | **SUB** | **H1** | **H2** | **H3** | **H1** | **H2** | **H3** | **H1** | **H2** | **H3** | **H1** | **H2** | **H3** | **H1** | **H2** | **H3** |
| CUR | SUB0 | 0.65 | 0.52 | 0.49 | 0.62 | 0.46 | 0.44 | 0.60 | 0.49 | 0.45 | 0.62 | 0.48 | 0.41 | 0.63 | 0.54 | 0.47 |
|  | SUB25 | 0.64 | 0.51 | 0.49 | 0.60 | 0.44 | 0.43 | 0.60 | 0.48 | 0.44 | 0.62 | 0.47 | 0.41 | 0.64 | 0.54 | 0.47 |
|  | SUB50 | 0.63 | 0.52 | 0.50 | 0.58 | 0.43 | 0.43 | 0.60 | 0.48 | 0.44 | 0.62 | 0.47 | 0.41 | 0.65 | 0.55 | 0.48 |
|  | SUB75 | 0.63 | 0.52 | 0.50 | 0.57 | 0.43 | 0.43 | 0.60 | 0.48 | 0.45 | 0.62 | 0.47 | 0.41 | 0.65 | 0.55 | 0.48 |
|  | SUB100 | 0.63 | 0.52 | 0.50 | 0.57 | 0.43 | 0.43 | 0.59 | 0.48 | 0.45 | 0.62 | 0.47 | 0.41 | 0.65 | 0.55 | 0.48 |
| SSP1-2.6 | SUB0 | 0.70 | 0.53 | 0.48 | 0.65 | 0.47 | 0.44 | 0.61 | 0.50 | 0.44 | 0.62 | 0.53 | 0.47 | 0.66 | 0.54 | 0.46 |
|  | SUB25 | 0.69 | 0.52 | 0.49 | 0.62 | 0.45 | 0.42 | 0.62 | 0.49 | 0.44 | 0.62 | 0.53 | 0.48 | 0.68 | 0.53 | 0.46 |
|  | SUB50 | 0.69 | 0.53 | 0.50 | 0.60 | 0.44 | 0.42 | 0.62 | 0.49 | 0.44 | 0.62 | 0.54 | 0.49 | 0.69 | 0.54 | 0.46 |
|  | SUB75 | 0.68 | 0.53 | 0.50 | 0.59 | 0.43 | 0.43 | 0.62 | 0.49 | 0.44 | 0.62 | 0.55 | 0.50 | 0.70 | 0.54 | 0.46 |
|  | SUB100 | 0.68 | 0.53 | 0.50 | 0.59 | 0.43 | 0.43 | 0.62 | 0.49 | 0.44 | 0.62 | 0.55 | 0.50 | 0.70 | 0.54 | 0.46 |
| SSP5-8.5 | SUB0 | 0.67 | 0.37 | 0.33 | 0.63 | 0.40 | 0.37 | 0.60 | 0.44 | 0.37 | 0.62 | 0.55 | 0.60 | 0.64 | 0.41 | 0.37 |
|  | SUB25 | 0.66 | 0.20 | 0.08 | 0.61 | 0.34 | 0.34 | 0.60 | 0.42 | 0.35 | 0.61 | 0.92 | 0.52 | 0.66 | 0.34 | 0.31 |
|  | SUB50 | 0.66 | 0.45 | 0.76 | 0.58 | 0.29 | 0.31 | 0.60 | 0.41 | 0.35 | 0.61 | 0.75 | 0.50 | 0.67 | 0.25 | 0.21 |
|  | SUB75 | 0.65 | 10.29 | 0.59 | 0.57 | 0.26 | 0.29 | 0.60 | 0.40 | 0.34 | 0.61 | 0.71 | 0.50 | 0.68 | 0.22 | 0.12 |
|  | SUB100 | 0.65 | 11.31 | 0.57 | 0.56 | 0.26 | 0.28 | 0.60 | 0.40 | 0.34 | 0.61 | 0.71 | 0.50 | 0.68 | 0.21 | 0.10 |

Table 11.  Parametric and smooth term estimates from the generalized additive model (GAM). The table presents estimated coefficients for the intercept and F-statistics with associated p-values for smooth terms. Significance levels: ***p < 0.001, **p < 0.01, *p < 0.05.*

| **Term** | **Estimate / F** | **p-value** | **Significance** |
| --- | --- | --- | --- |
| (Intercept) | 0.49825 | <2e-16 | *** |
| s(total_CM):ClimateCUR | F = 10.50 | 0.000355 | *** |
| s(total_CM):ClimateSSP26 | F = 12.98 | 5.04e-05 | *** |
| s(total_CM):ClimateSSP85 | F = 101.72 | <2e-16 | *** |
| s(duration):ClimateCUR | F = 9.90 | 0.001905 | ** |
| s(duration):ClimateSSP26 | F = 7.36 | 0.000542 | *** |
| s(duration):ClimateSSP85 | F = 62.75 | <2e-16 | *** |

**Note:**
*The model explained 80.3% of the deviance (adjusted R² = 0.794, n = 211)*

Table 12:  Parametric and smooth term estimates from the generalized additive model (GAM). The table shows the intercept estimate and F-statistics with corresponding p-values for each smooth term. Significance levels: ***p < 0.001, **p < 0.01, *p < 0.05, ·p < 0.1.*

| **Term** | **Estimate / F** | **p-value** | **Significance** |
| --- | --- | --- | --- |
| (Intercept) | 0.52124 | <2e-16 | *** |
| s(total_emission):ClimateCUR | F = 3.996 | 0.063892 | . |
| s(total_emission):ClimateSSP26 | F = 5.104 | 0.024945 | * |
| s(total_emission):ClimateSSP85 | F = 37.397 | <2e-16 | *** |
| s(total_CM):ClimateCUR | F = 7.748 | 0.000949 | ** |
| s(total_CM):ClimateSSP26 | F = 8.870 | 0.000912 | *** |
| s(total_CM):ClimateSSP85 | F = 44.369 | <2e-16 | *** |

**Note:**
*The model explained 75.1% of the deviance (adjusted R² = 0.740, n = 211)*

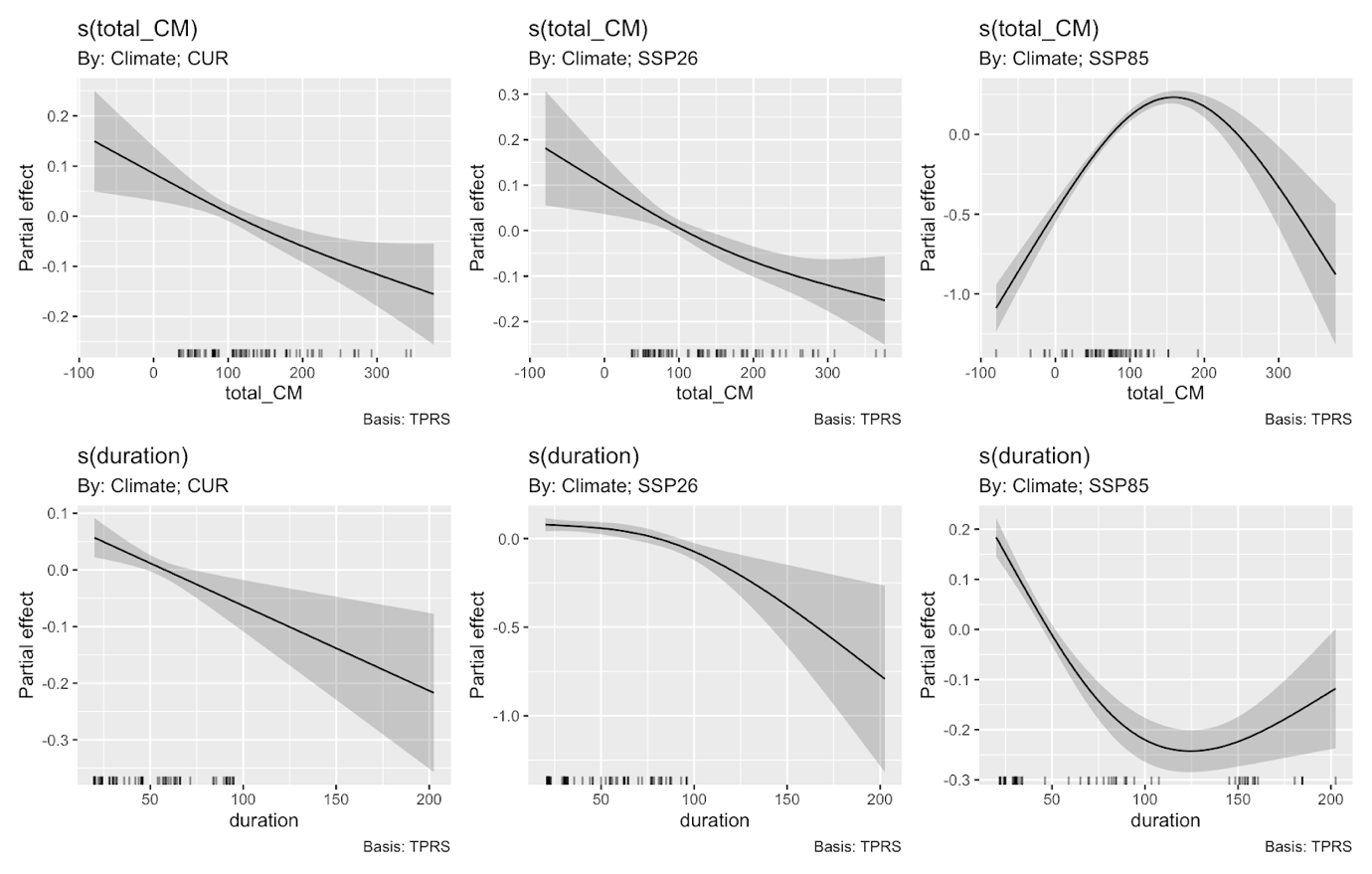

Figure 1: Partial effects from GAM predicting mitigation efficiency as a function of cumulative carbon balance (total_CM) and emission duration (duration), stratified by climate scenario (CUR, SSP1-2.6, SSP5-8.5). Each panel shows smoothed effects (s(x, k=3)) with 95% confidence intervals.

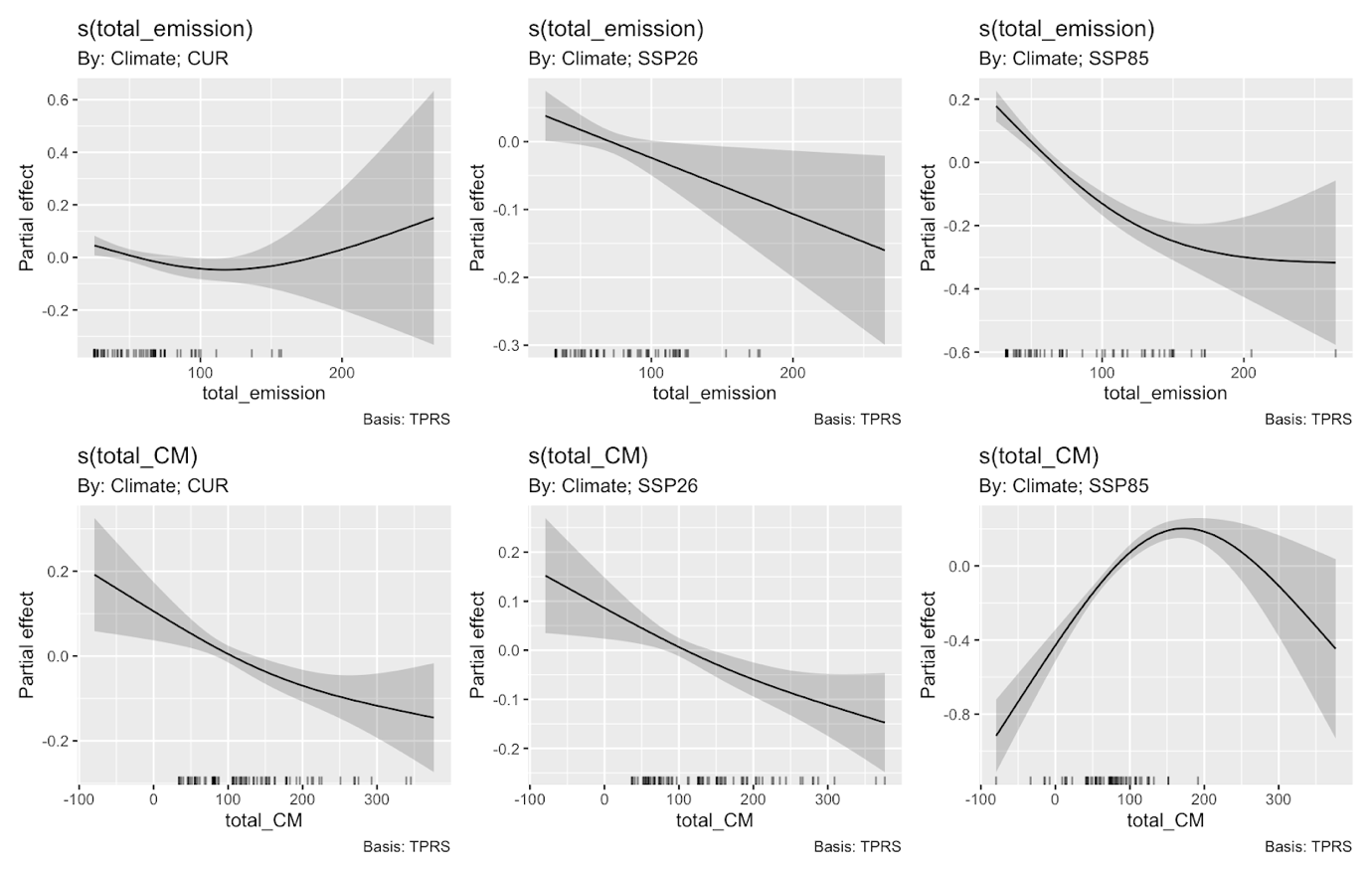

Figure 2: Partial effects from GAM predicting mitigation efficiency as a function of cumulative carbon balance (total_CM) and emission amplitude (total_emission), stratified by climate scenario (CUR, SSP1-2.6, SSP5-8.5). Each panel shows smoothed effects (s(x, k=3)) with 95% confidence intervals.

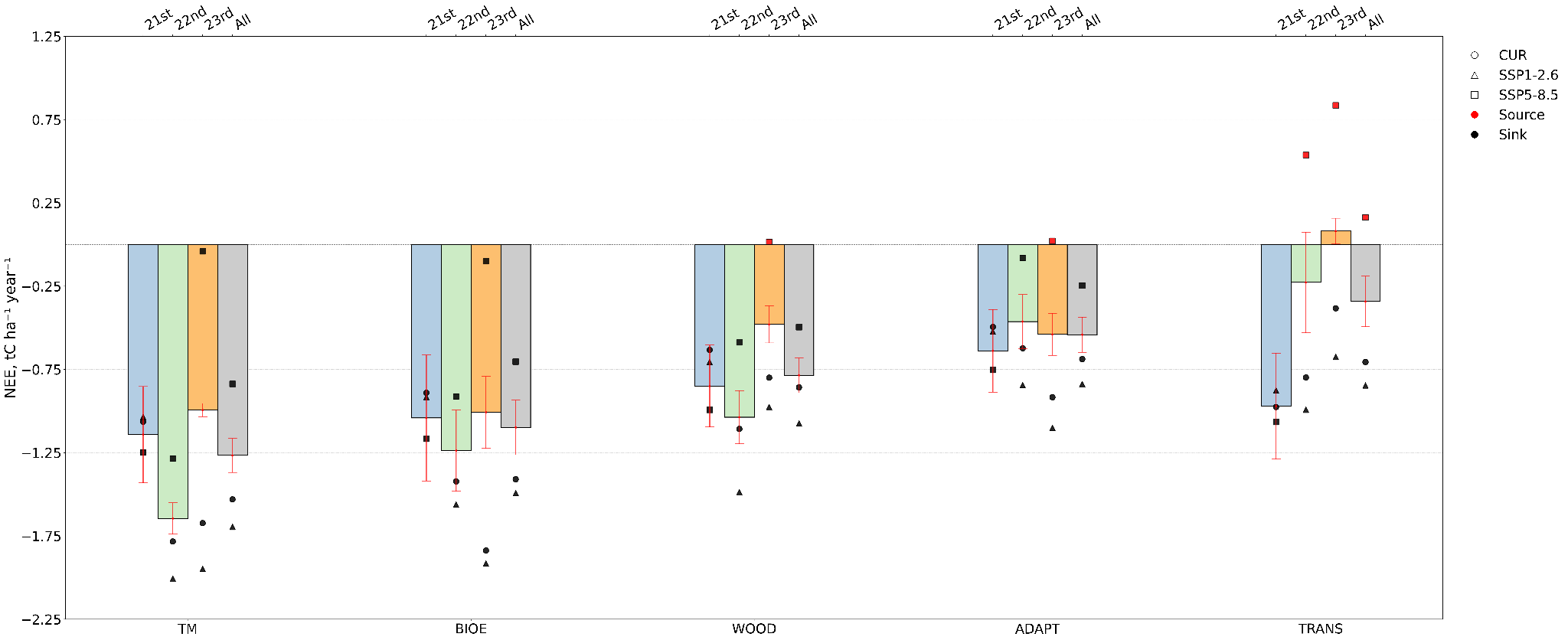

Figure 3: Mean NEE in tC ha^-1^ year^-1^ for five forest management strategies (TM, ADAPT, BIOE, TRANS, and WOOD), averaged across two climate scenarios (SSP1-2.6 and SSP5-8.5), shown for the 21st, 22nd, and 23rd centuries, as well as for the full period combined. Red error bars indicate the 95% confidence interval of NEE across climate scenarios. The forest ecosystem is a net C-sink (negative NEE) when markers are black and a net C-source when markers are red (positive NEE).

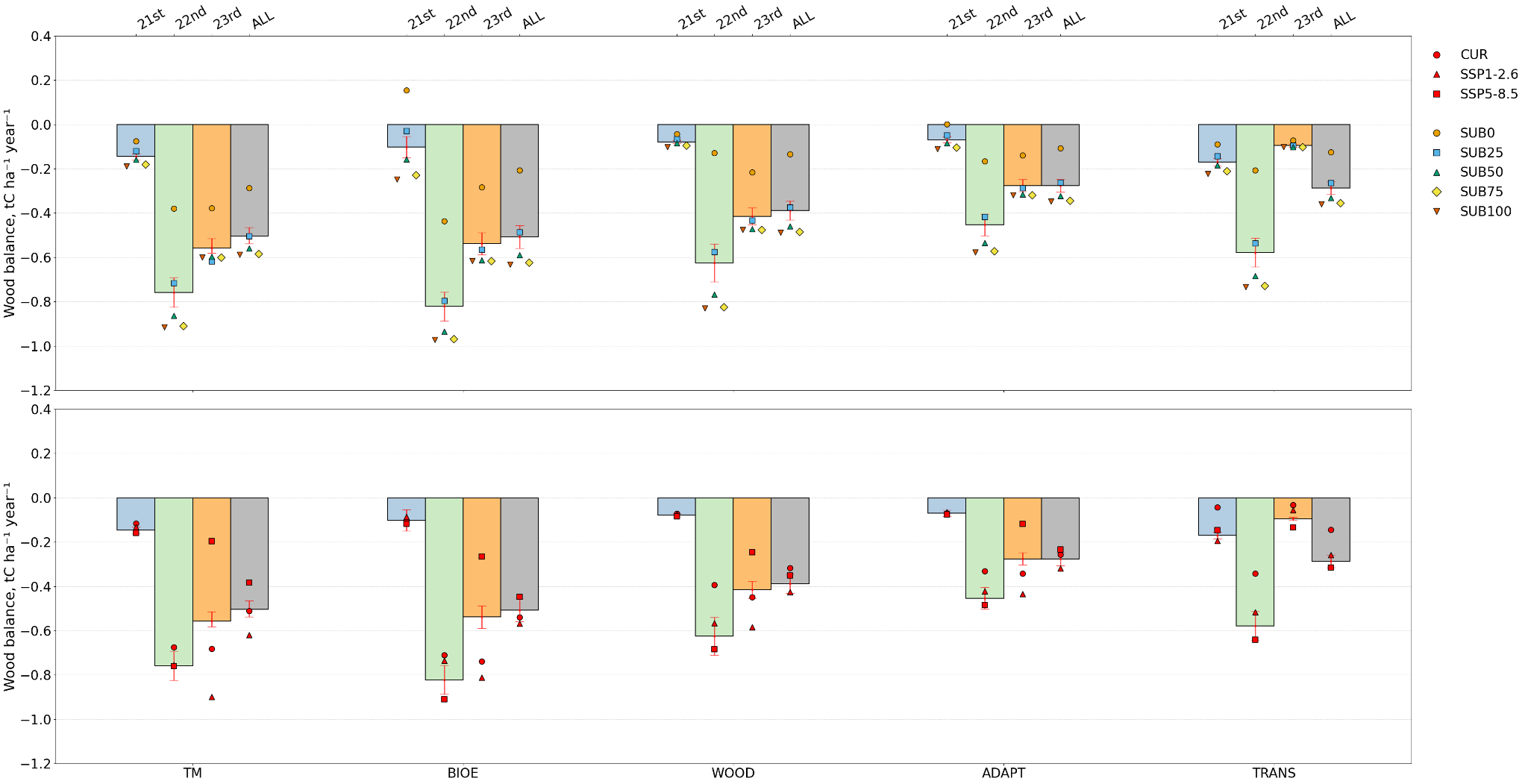

Figure 4: Mean wood balance (Wbalance), in tC ha^-1^ year^-1^ for five forest management strategies (TM, ADAPT, BIOE, TRANS, and WOOD), averaged across two climate scenarios (SSP1-2.6 and SSP5-8.5), shown for the 21st, 22nd, and 23rd centuries, as well as for the full period combined. Red error bars indicate the 95% confidence interval of Wbalance across climate scenarios. The upper panel shows Wbalance values by decarbonization pathway, while the lower panel shows Wbalance values by climate scenario.
